# Membrane estrogen receptor (GPER) and follicle-stimulating hormone receptor heteromeric complexes promote human ovarian follicle survival

**DOI:** 10.1101/2020.04.21.053348

**Authors:** Livio Casarini, Clara Lazzaretti, Elia Paradiso, Silvia Limoncella, Laura Riccetti, Samantha Sperduti, Beatrice Melli, Serena Marcozzi, Claudia Anzivino, Niamh S. Sayers, Jakub Czapinski, Giulia Brigante, Francesco Potì, Antonio La Marca, Francesco De Pascali, Eric Reiter, Angela Falbo, Jessica Daolio, Maria Teresa Villani, Monica Lispi, Giovanna Orlando, Francesca G. Klinger, Francesca Fanelli, Adolfo Rivero-Müller, Aylin C. Hanyaloglu, Manuela Simoni

## Abstract

Classically, follicle stimulating hormone receptor (FSHR) driven cAMP-mediated signaling boosts human ovarian follicle growth and would be essential for oocyte maturation. However, contradicting *in vitro* suggest a different view on physiological and clinical significance of FSHR-mediated cAMP signaling. We found that the G protein coupled estrogen receptor (GPER) heteromerizes with FSHR, reprogramming cAMP/death signals into proliferative stimuli fundamental for sustaining oocyte survival. In human granulosa cells, survival signals are effectively delivered upon equal expression levels of both receptors, while they are missing at high FSHR:GPER ratio, which negatively impacts follicle maturation and strongly correlates with FSH responsiveness of patients undergoing controlled ovarian stimulation. Consistent with high FSHR expression levels during follicular selection, cell viability is dramatically reduced in FSHR overexpressing cells due to preferential coupling to the Gαs protein/cAMP pathway. In contrast, FSHR/GPER heteromer formation resulted in FSH-triggered anti-apoptotic/proliferative signaling delivered via the Gβγ dimer while heteromer impairment or GPER-associated Gαs inhibitory protein complexes resulted in cell death. GPER-depleted granulosa cells have an amplified FSH-dependent decrease in cell viability and steroidogenesis, consistent with the requirement of estrogen signaling for successful oocyte growth. Therefore, our findings indicate how oocyte maturation depends on the capability of GPER to shape FSHR selective signals, indicating hormone receptor heteromers may be a marker of cell proliferation.

**One Sentence Summary:** FSHR/GPER heteromers block cAMP-dependent selection of ovarian follicles and target tumor growth and poor FSH-response in women.

## Introduction

Ovarian follicular growth and dominance in women of reproductive age is a physiological example of how a tightly regulated equilibrium between active cell proliferation and apoptosis results in the selection of a single dominant follicle at the expense of all others. Key players of this game are sex-hormones, follitropin (FSH) and 17β-estradiol (E_2_), which stimulate cell viability and proliferative signals in the gonads and certain tumor cells^1,2^ Sex-hormone receptors are druggable targets in fertility and cancer treatment to control cell death and survival.

The FSH-receptor (FSHR) stimulates Gαs protein-dependent cAMP/PKA activation, resulting in cAMP-response element binding protein (CREB) phosphorylation and steroidogenic activity, necessary to produce estrogens that, in turn, are well-known stimulators of growth^3^. However, a pro-apoptotic role of FSH has also been proposed^4,5^ and, intriguingly, prolonged FSHR overexpression^6^ or accumulation of high intracellular cAMP levels^7,8^ are a prerequisite for both steroid synthesis and cell death^9^ The FSH-related pro-apoptotic activity occurs when high FSHR expression is induced^6^, providing a plausible reason why no consistent steroidogenic cell lines permanently overexpressing the FSHR exist so far^10,11^. Other FSHR functions have been reported to be mediated by Gαi and Gαq proteins, the Gβγ dimer^12,13^ and other molecules inducing proliferative signals under low FSHR density in the cell membrane^14^ FSHR-mediated activation of protein kinase B (AKT) occurs downstream of G protein activation^15–17^, and results in anti-apoptotic and proliferative activity in FSHR-expressing ovarian^18^ and cancer^19^ cells.

Estrogens activate two nuclear receptors, ERα and ERβ, and a membrane G protein-coupled receptor (GPCR), named GPER, mediating rapid E_2_-induced intracellular responses such as calcium ion (Ca^2+^) mobilization^20^ and found in ovarian tissues throughout the follicular phase^21^. Although GPER is coupled to Gαs, it is unable to trigger intracellular cAMP accumulation in response to E_2_ in a number of cell models^22^ This is due to the interaction of GPER with both the membrane-associated guanylate kinases (MAGUK) and protein kinase A-anchoring protein 5 (AKAP5), constitutively inhibiting cAMP in a Gαi/o-independent manner^22^ GPER mediates survival/proliferation signals through phosphoinositide 3 (PI3K)/phospho-AKT (pAKT) and phospho-extracellular-regulated kinases 1 and 2 (pERK1/2) activation^23^, as well as Gβγ-dependent mechanisms upregulating proto-oncogenes^24^.

Ovarian granulosa cells express both FSHR and GPER^25^ modulating a network of proliferative signals^26,27^ fundamental for regulating gametogenesis^21^. Consistent with its impact on cell growth, FSHR expression was found in pathological contexts characterized by uncontrolled cell proliferation, such as tumors^28^ or endometriosis^29^, suggesting that it could be a target for still outstanding anti-cancer therapies^30^. Similarly, GPER expression has been described in several tumor cells^31^, including breast, endometrium and ovary^32^, where it may cooperate with FSHR in inducing uncontrolled cell proliferation^27,33^.

Physiologically, high FSHR expression occurs transitorily in the ovarian granulosa cells, decreasing once a single dominant follicle is selected^34^. We reasoned that the opposing nature of FSHR activity, proliferative and pro-apoptotic, may rely on the cooperation with other factors modulating FSHR signaling. In particular, as described for other structurally similar GPCRs^35–38^, FSHR signaling may be regulated by forming heteromers with other receptors^10^. Here, we demonstrate that GPER and FSHR interactions reprogram FSHR-related death into life signals, upregulating the viability of FSHR/GPER expressing cells including human ovarian granulosa cells. Together, our data provides a mechanistic model for dominant follicle selection in humans and a novel therapeutic target for poor FSH-responder women undergoing controlled ovarian stimulation.

## Results

### FSHR and GPER co-expression is linked to ovarian follicle maturation: in vivo, pharmacological relevance in an assisted reproduction setting

We first evaluated the presence of both FSHR and GPER in granulosa cells as demonstrated by immunostaining of human ovarian follicle tissue sections (Fig. 1A, B) after antibody validation (data file S1). This was confirmed by Western blotting of human granulosa and transfected HEK293 cell lysates, in the presence and in the absence of *GPER* mRNA depletion by siRNA (Fig. 1C), where the anti-FSHR or anti-GPER antibodies produced a signal in cells expressing only these receptors (Fig. 1C).

**Fig. 1.**
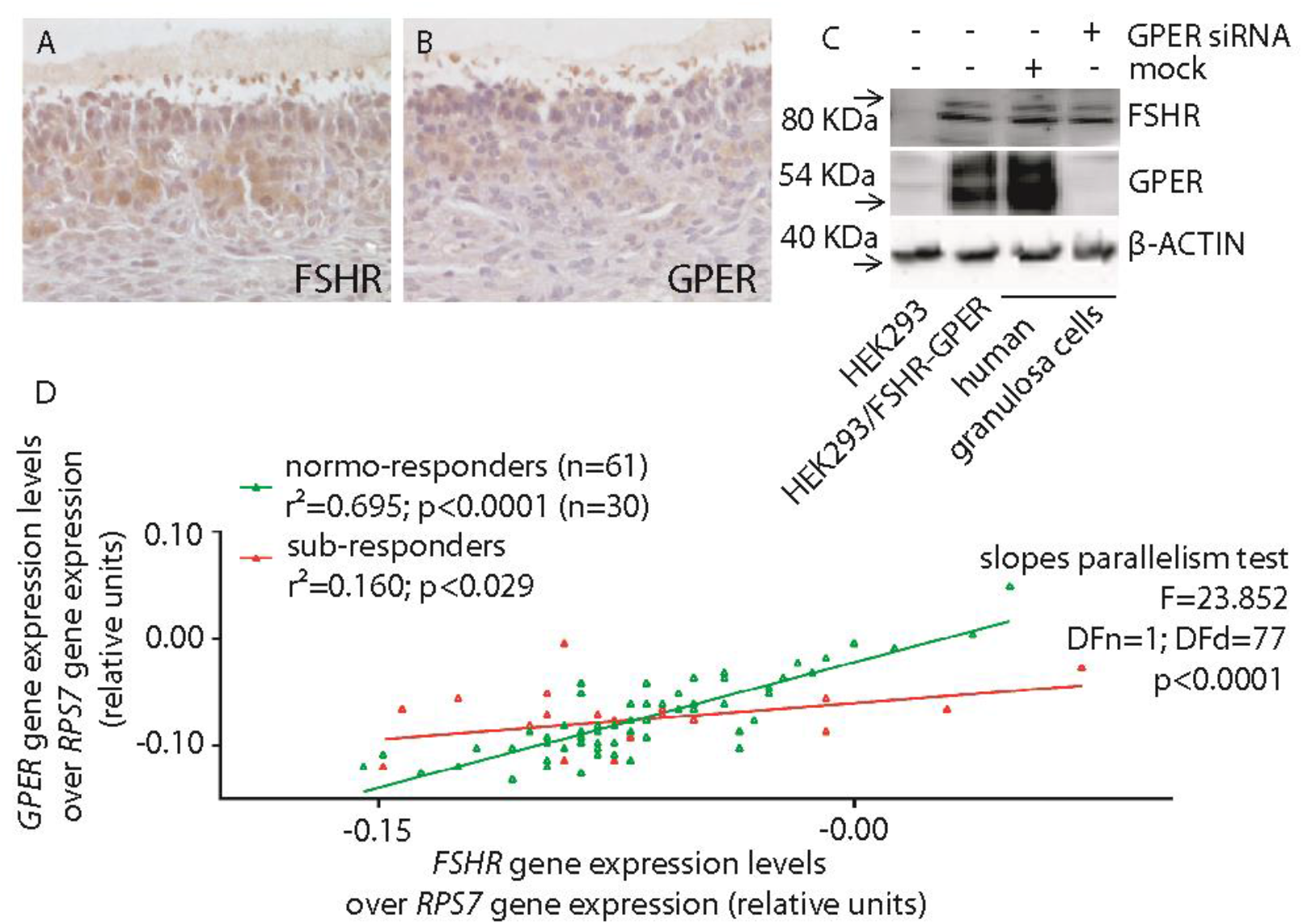
Link between FSHR-GPER co-expression and follicular growth response to FSH. (**A** and **B**) Representative determination of FSHR and GPER expression in granulosa cells at the antral follicular stage, by immunohistochemistry. Ovarian sections were treated by specific anti-FSHR or -GPER primary antibodies, over hematoxylin background staining. (**C**) Western blotting demonstrating the presence of both FSHR and GPER in primary human granulosa cell lysates and the inhibition of expression of GPER by 48-h siRNA. HEK293 transiently transfected with FSHR- and GPER-encoding plasmids were used as controls. The efficacy of GPER siRNA is shown and compared with control siRNA (mock)-treated granulosa cells. β-ACTIN was used as a loading control. (**D**) Correlation between *FSHR* and *GPER* gene expression levels in granulosa cells collected from donor normo-(n=61) and sub-responder (n=30) women undergoing FSH stimulation for assisted reproduction. Each patient is represented by a point and mRNA levels were measured by real-time PCR, normalized over the *RPS7* housekeeping gene and interpolated by linear regression.

To investigate if FSHR-GPER co-expression may have physiological and clinical relevance, *FSHR*/*GPER* mRNA expression levels were evaluated in granulosa cells collected from ovarian follicular fluids of women undergoing FSH stimulation for assisted reproduction techniques (ART). This data was then matched with clinical parameters indicative of *in vivo* proliferation. A group of 91 women were subdivided in two groups; “normo-responders” were defined as women providing >4 oocytes upon controlled ovarian stimulation, while ≤4 collected oocytes defined as “sub-responders”. Granulosa cells from donor women were cultured one week to recover basal *FSHR* and *GPER* expression, and receptor mRNA levels quantified, plotted in a X-Y graph and interpolated by non-linear regression. In normo-responder women, *FSHR* expression increased linearly together with *GPER* transcripts, suggesting that a certain amount of these two receptors should be maintained within a certain ratio for inhibiting pro-apoptotic signals and dampening follicular growth (Fig. 1D; data file S2). The functional importance of such a ratio is supported by the finding that in FSH sub-responders, low oocyte yield is correlated to higher *FSHR* over *GPER* expression, suggesting this may impact cell survival.

### FSH and cAMP pro-apoptotic potential is associated with FSHR expression levels and is counteracted by E_2_

We evaluated the FSHR-related potential of activating steroidogenic and pro-apoptotic signals, likely associated with cAMP production^3,7^ Intracellular accumulation of cAMP depends on FSHR-Gαs protein coupling, which constitutively increases together with receptor expression levels^39^ This was demonstrated by employing Gαs- and FSHR-tagged BRET biosensors, which resulted in left-shifted BRET saturation curves at the maximal concentration of receptor-encoding plasmid used to transfect HEK293 (HEK293/FSHR) cells. This indicates a greater receptor-G protein association affinity is achieved when FSHR is highly expressed (Fig. 2A; data file S3). The receptor expression-dependent association with Gαs was not observed with other G proteins reported to be FSHR intracellular interactors, such as Gαi and Gαq (Fig. 2B-E; data file S4; data file S5). While Gαs likely competes with other interactors for binding a low number of FSHRs, the potential of mediating both basal and FSH-induced cAMP activation was FSHR concentration-dependent (Fig. 2F). Increases in intracellular cAMP is deleterious for the viability of FSHR-expressing cells^6^. Indeed, treatment of either HEK293/FSHR or human primary granulosa cells with 8-br-cAMP, induces reduction of cell viability in a concentration-dependent manner (Fig. 2G, H). Interestingly, the cAMP analog-induced decrease of granulosa cell viability was inhibited by 50 pg/ml E_2_ (Fig. 2H), suggesting possible cross-talk between opposing gonadotropin- and estrogen-mediated intracellular actions. Indeed, in the absence of E_2_, FSH-induced a decrease of HEK293/FSHR cell viability (Fig. 2I, J).

**Fig. 2.**
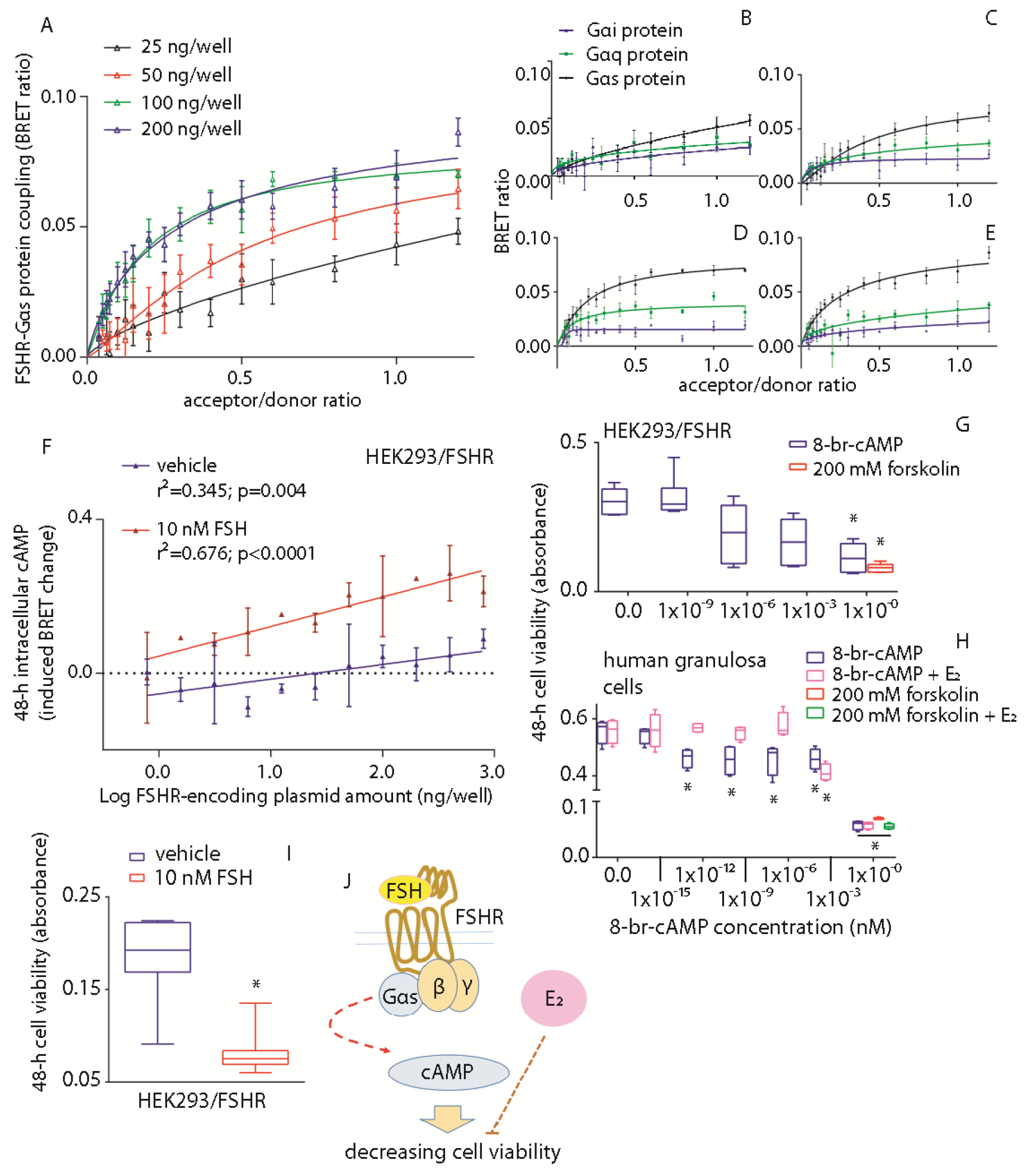
Excessive FSHR expression levels impact negatively HEK293 cell viability. (**A**) Association of FSHR to the Gαs protein increases together with receptor expression levels. Coupling experiments were performed using HEK293 cells co-expressing the FSHR/rluc-(donor) and the Gαs protein/venus-tagged BRET biosensor. Data were interpolated by non-linear regression and compared by Kruskal Wallis test and Dunn’s post-test (100 and 200 ng/well of FSHR-encoding plasmid *versus* the 25 ng/well condition; p=0.0014; means ± SEM; n=5). (**B-E**) Comparison of constitutive FSHR coupling to Gαs, Gαq and Gαi with increasing receptor expression levels, in HEK293 cells transfected using 25 (**B**), 50 (**C**), 100 (**D**) and 200 (**E**) ng/well of FSHR/rluc-encoding plasmid. Receptor level-dependent increase in affinity occurs with Gαs but not with the other proteins (Kruskal Wallis test and Dunn’s post-test; p<0.0001; means ± SEM; n=5). (**F**) Basal and FSH-induced intracellular cAMP levels with increasing FSHR expression levels. Data were represented as means ± SEM and interpolated by linear regression (n=4). (**G**) Colorimetric assay reveals the relationship between increasing intracellular cAMP concentration and decreasing of HEK293/FSHR cell viability. Cells were treated with increasing concentrations of 8-br-cAMP, while 200 mM forskolin-treated cells were positive controls. Data were represented by box and whiskers plots (*=significantly different *versus* 0.0 8-br-cAMP-treated samples; Kruskal Wallis test and Dunn’s post-test; p=0.0019). (**H**) In human primary granulosa lutein cells, 50 pg/ml E_2_ co-treatment inhibits the decrease in cell viability induced by the cAMP analog (*=significantly different *versus* 0.0 8-br-cAMP-treated samples; Kruskal Wallis test and Dunn’s post-test; n=8). (**I**) Cell viability is lower in HEK293/FSHR transfected cells stimulated with FSH compared to vehicle (Mann-Whitney’s *U*-test; p=0.0003; n=8). (**J**) Summary: FSHR coupling to the Gαs protein/cAMP-pathway, occurring at high receptor expression levels and in the absence of E_2_, decreases cell viability. This action is counteracted by estrogen.

### FSHR and GPER form heteromeric complexes at the cell membrane

In order to define the mechanism linking FSH and GPER/estrogen-mediated intracellular networks, we employed distinct approaches to examine the formation of possible heterodimers/oligomers involving FSHR and GPER at the cell membrane. Investigation at the atomic level relied on a protein-protein docking-based approach, the FiPD-based approach^40,41^ applied to the structural models of the two receptors. The two independent docking runs by using FSHR as a target and GPER as a probe and *vice versa* converged on the same predicted architecture of the heterodimer (Fig. 3A), ranked among the best ten out of 4000 solutions and characterized by good membrane topology. Remarkably, when using FSHR as a target, the predicted docking solution was the best in score (i.e. rank #1) out of 4000, belonged to the most populated solution cluster, and showed a good membrane topology (MemTop) score (0.578) (see Methods). The FSHR-GPER interface in the predicted heterodimer is characterized by contacts between H6 and H7 from FSHR and H7 and H6 from GPER, respectively (Fig. 3A).

**Fig. 3.**
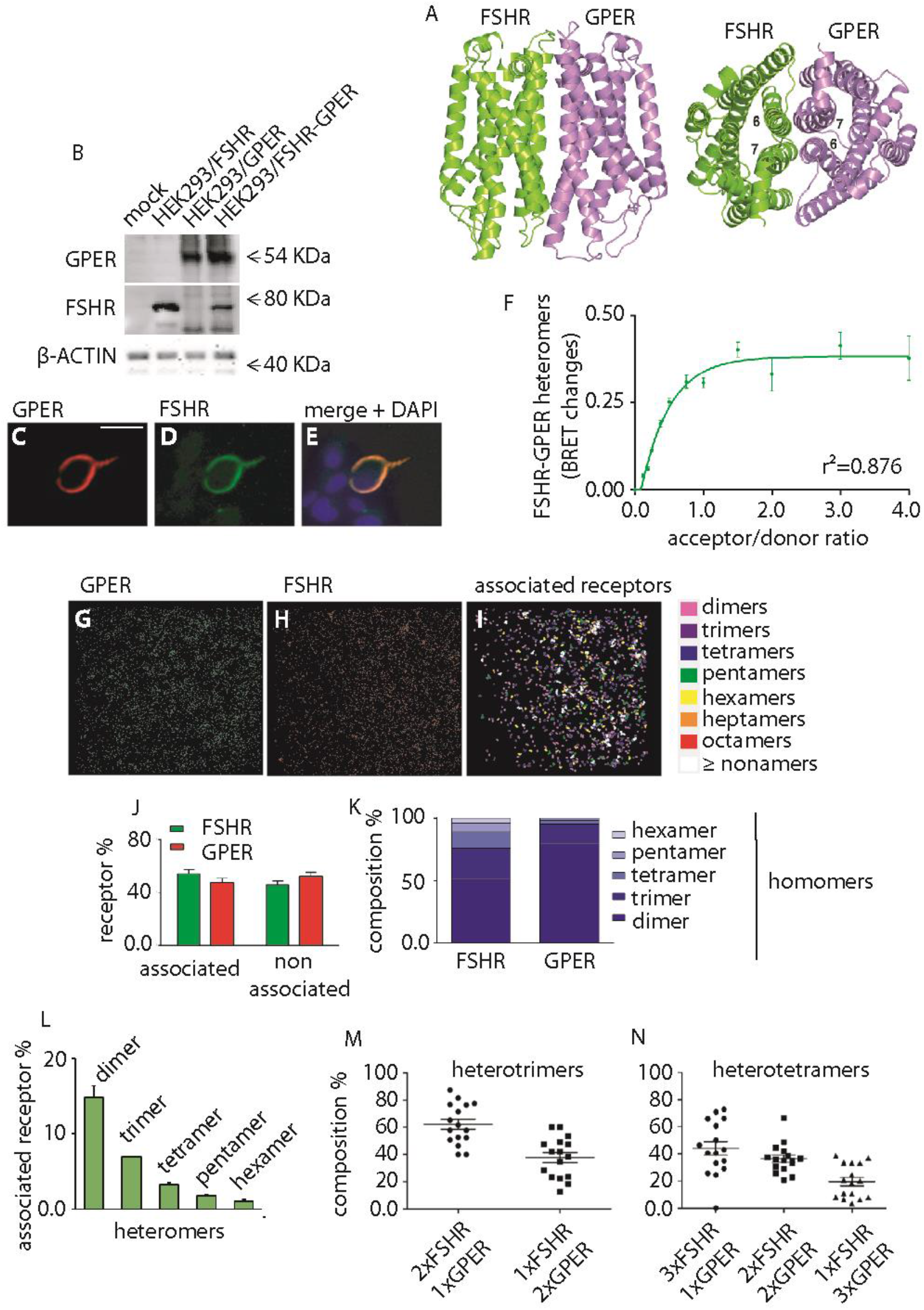
The FSHR forms heteromers with GPER. (**A**) Predicted structural model of the heterodimer between FSHR (green) and GPER1 (violet) seen in directions perpendicular (left) and parallel (right) to the bundle main axis. In this dimer, H6 of FSHR interacts with H7 of GPER1 and H6 of GPER1 interacts with H7 of FSHR. (**B**) Control Western blotting for FSHR and GPER transient expression and co-expression in HEK293 cells. β-ACTIN served as normalizer. (**C-E**) Representative evaluation of GPER and FSHR co-localization by immunofluorescence, in HEK293 cells transiently transfected with FSHR and GPER. A specific primary antibody was used for GPER binding before cell treatment by a TRITC-labelled secondary antibody, while nuclei were blue-stained by DAPI. FSHR is identified by the venus tag light emission. Bar = 25 μm. (**F**) Formation of FSHR/rluc- and GPER/venus-tagged heteromers in transfected HEK293 cells. BRET ratio values resulting from molecular interactions were represented as means ± SEM. Specific association is indicated by data interpolation using non-linear regression, which results in the logarithmic curve (n=4). Unspecific binding is provided as supplemental data S3. (**G, H**) Confirmation of FSHR-GPER heteromers visualized by photo-activated localization microscopy with photo-activatable dyes (PD-PALM) in HEK293 cells. **(I)** Representative PD-PALM images and associated molecule heat maps from cells expressing FLAG-GPER and HA-FSHR. Images are reconstructed from 7-7 μm^2^ areas of plasma membrane after x-y coordinate localization using QuickPALM followed by a 50 nm radius neighborhood analysis of receptor molecules. (**J**) Quantitative analysis of associated versus non-associated FSHR and GPER homomers using single channel PD-PALM, Mean ± SEM, n = 5. (**K**) Quantitative evaluation of the types of FSHR and GPER homomers. Data is expressed as percentage of total associated complexes. (**L**) Quantitative analysis of heteromeric assemblies between FSHR and GPER using dual channel PD-PALM reveals the prevalence of heterodimeric formations, Mean ± SEM, n = 8 cells. (**M, N**) Analysis of individual protomer composition within heterotrimers (**M**) and heterotetramers (**N**), demonstrates the preferential occurrence of FSHR protomers *versus* GPER protomers within these individual multimers, Mean ± SEM, n = 8.

Western blotting and immunofluorescent staining of HEK293/FSHR-GPER cells confirmed expression of both receptors in cell lysates (Fig. 3B) and their co-localization at the cell surface (Fig. 3C-E). No signals were detected in GPER- and FSHR-negative cells (Fig. 3E). The physical interaction between the two receptors was demonstrated by BRET. In transfected HEK293 cells, transiently expressing both FSHR-rluc and venus-tagged GPER (GPER/rluc) biosensors, BRET signal logarithmically increases together with the acceptor concentration, indicating specific heteromeric interactions between the two receptors (Fig. 3F; data file S6; data file S7). Further evidence of FSHR-GPER heteromer assembly at the plasma membrane was provided by photo-activated localization microscopy with photoactivatable dyes (PD-PALM) (Fig. 3G-I), a super-resolution imaging approach we have previously employed to quantitate protomer composition within asymmetric heteromer complexes between LHR mutants and between FSHR and LHR^38,42^ HEK 293 cells expressing HA-tagged FSHR and FLAG-tagged GPER individually, or together, were labeled with CAGE500-conjugated anti-HA and CAGE552-conjugated anti-FLAG antibodies and fixed for PD-PALM imaging (Fig. 3G-I). FSHR and GPER existed as pre-existing monomers and homomers in equivalent amounts (Fig. 3J), although while FSHR exhibited a range of homomeric complexes, GPER-GPER associations were primarily dimeric (Fig. 3K). GPER-FSHR heteromers at the plasma membrane were also observed in cells co-expressing these receptors, consistent with the BRET data. The primary heteromer population represented heterodimers but also exhibited a range of heterooligomeric complexes (Fig. 3L). Further analysis of the protomer identity within individual hetero-oligomeric complexes revealed such complexes favored formation of FSHR dominant hetero-oligomers (Fig. 3M, N), suggesting potential asymmetry in these low order heterooligomers. Overall, this data demonstrates that FSHR and GPER form distinct heteromeric assemblies at the plasma membrane and suggests the existence of a novel molecular mechanism modulating the combined action of FSH and estrogen.

### FSH stimulation of FSHR-GPER heteromers promotes cell viability via inhibition of cAMP signaling

The function of GPER and its cooperation with FSHR in modulating cell viability was next investigated. HEK293/GPER expressing cells displayed rapid intracellular Ca^2+^ increase (Fig. 4A) occurring immediately upon 10 pg/ml estradiol addition, demonstrating expression of functional GPER^20^. BRET measurements also revealed that GPER can couple to Gαs (Fig. 4B), although the receptor is unable to mediate cAMP production upon ligand binding (Fig. 4C), consistent with previous reports demonstrating the association of GPER with a cAMP/PKA inhibitory complex assembled by MAGUK and AKAP5 proteins^22^ Intracellular cAMP levels were maximally stimulated upon FSH treatment of HEK293/FSHR cells but, interestingly, not in HEK293 cells co-expressing FSHR and GPER (Fig. 4C). These data indicate that the presence of GPER inhibits the increase of intracellular cAMP, independent of the presence or absence of E_2_, in FSHR-expressing cells. This effect is specifically targeted to FSHR, since LH treatment of LHCGR-GPER co-expressing HEK293 cells induces an increase in intracellular cAMP (data file S8). Moreover, GPER-dependent inhibition of FSH-induced cAMP signaling occurs even under blockade of the nuclear estrogen receptor by fulvestrant (Fig. 4D), thus excluding its involvement. The ability of GPER to inhibit FSH-mediated cAMP signaling was confirmed by measurement of FSH-induced CREB phosphorylation, a downstream cAMP-dependent event (Fig. 4E).

**Fig. 4.**
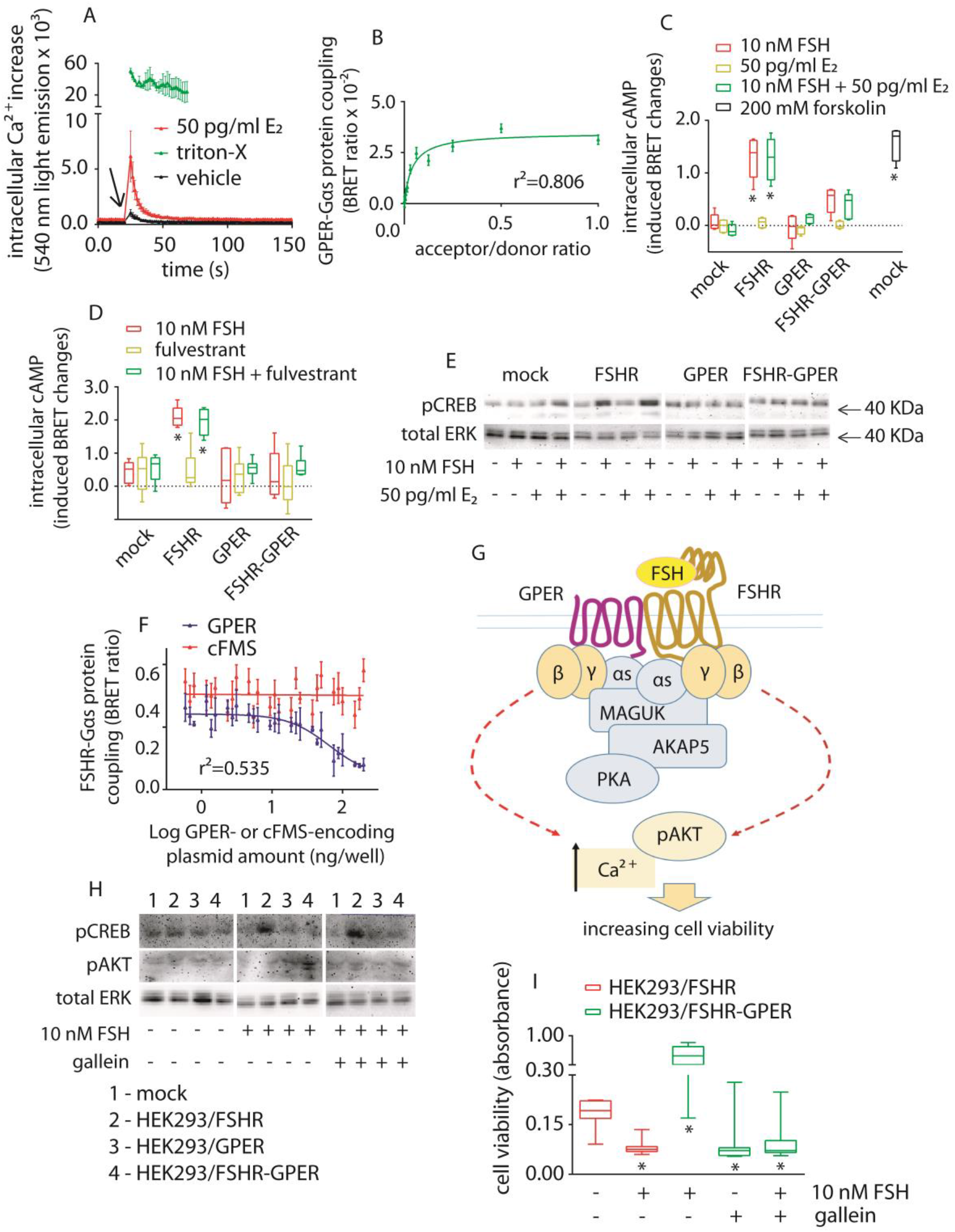
FSHR-GPER crosstalk promotes FSH-stimulated cell viability via Gβγ dimers. (A) GPER transiently expressed in HEK 293 cells simulates E_2_-induced intracellular Ca^2+^ increase (Kruskal-Wallis test; p<0.0001; n=8; means ± SEM). Signals were captured by BRET over 150 s, in the presence of the calcium-biosensor. PBS (vehicle)-treated cells provided basal levels. Cells lysed by triton-X were positive controls and only a 43-s time window is representatively shown. Compounds were added at the 21-s time-point. (B) GPER/rluc-coupling to the Gαs protein/venus-tagged was demonstrated by BRET. Values are means ± SEM and the logarithmic curve was obtained after interpolation using non-linear regression (n=8). (C) 10 nM FSH induced cAMP increase in HEK293 cells expressing either one or both FSHR and GPER. 50 pg/ml E_2_ were added as indicated, while mock-transfected and forskolin-treated cells served as basal and positive control, respectively. cAMP values are indicated as induced BRET changes over vehicle-treated mocks (*=significantly different versus FSH-treated mocks; two-way ANOVA with Dunnett’s correction for multiple tests; p<0.0001; n=5; means ± SEM). (D) Evaluation of 10 nM FSH-induced intracellular cAMP increase, in the presence and in the absence of the nuclear estrogen receptor blockade by fulvestrant. *=significantly different versus FSH-treated mocks; two-way ANOVA with Dunnett’s correction for multiple tests; p<0.0001; n=5; means ± SEM. (E) Representative Western blotting analysis of the cAMP-dependent pCREB activation, in HEK293 cells expressing either one or both FSHR and GPER. 50 pg/ml E_2_ were added where indicated and total ERK served as normalizer. (F) Decrease of BRET signal indicating a reduced FSHR/rluc-Gαs protein/venus interaction along with increasing untagged GPER expression levels (n=3; means ± SEM). The receptor-tyrosine kinase colonystimulating factor-1 receptor (cFMS) was the negative control. (G) Proposed model of the FSHR/cAMP signaling blockade by the GPER-associated inhibitory machinery. The MAGUK/AKAP5 complex might physically interact with the Gαs protein coupled to FSHR upon heteromer formation, resulting in cAMP signaling blockade. At this point we hypothesized that FSH-dependent intracellular signals modulating cell viability could be activated by the βγ dimer. (H) Evaluation of Gαs protein-dependence of pCREB and βγ dimer-dependence of 15-min pAKT activation, in HEK293 cells expressing either one or both FSHR and GPER, respectively. The βγ dimer inhibitor gallein was used where indicated and total ERK was the normalizer. (I) Decreasing of 10 nM FSH-induced cell viability in FSHR-GPER-co-expressing HEK293 cells due to βγ dimer-blockade by gallein. Results were compared with cell viability data from gallein-untreated cells and HEK293/FSHR cells of Fig 2I (*=different versus mock; Kruskal-Wallis test; p<0.0001; n=8; means ± SEM).

To determine if FSHR and Gαs basal coupling was altered in cells co-expressing GPER, FSHR-Gαs coupling was measured via BRET. Increasing levels of GPER decreased BRET signals between FSHR and Gαs (Fig. 4F; data file S8). While it is unknown whether the decay of the BRET signal corresponds to an uncoupling or structural rearrangement of the FSHR-Gαs complex, it is indicative of a physical perturbation of such complex by GPER. We hypothesized that the GPER/MAGUK/AKAP5 complex associates with FSHR upon heteromer formation perturbing FSHR-Gαs pre-coupling. This disables the ability of FSH to increase cAMP, while not affecting Gβγ-dependent signaling by FSH (Fig. 4G). This hypothesis was explored by measuring Gβγ-dependent AKT phosphorylation by FSHR-GPER heteromers, in response to FSH (Fig. 4H). Acute FSH treatment (15 min) did not induce pAKT activation in HEK293/FSHR cells, which requires chronic FSH stimulation to detect activation^43^. In contrast, pAKT activation was increased in FSH-treated HEK293/FSHR-GPER cells and was inhibited by the selective Gβγ inhibitor gallein. GPER also increased cell viability following FSH treatment in FSHR-GPER cells (Fig. 4I), compared to HEK293/FSHR cells (Fig. 2I), which critically was also inhibited by gallein. Taken together, these results demonstrate that GPER is capable of downregulating FSHR/Gαs/cAMP to increase cell viability via a Gβγ-dependent mechanism.

### Disruption of FSHR-GPER heteromer formation results in FSH-induced decrease in cell viability

To demonstrate that, in cells co-expressing FSHR and GPER, FSH/FSHR-mediated increase in cAMP and cell viability was directly due to the FSHR-GPER heteromer formation, a mutant GPER (GPERmut) was created that would not form heteromers with FSHR. To this purpose, the predicted structural model of the heterodimer was exploited to drive site-directed mutagenesis. All the H6 and H7 non-glycine and non-alanine amino acids facing FSHR were replaced by alanines, leading to a GPER form mutated in fifteen positions distributed along the whole length of the two helices, the last amino acid being the beginning of H8 (Fig. 5A). GPERmut was still able to activate calcium signaling (Fig. 5B) and co-expressed with FSHR at the cell membrane (Fig. 5C-E). Despite exhibiting ability to activate calcium signaling, the GPERmut was unable to form heteromers with FSHR as demonstrated by the lack of a saturated BRET signal (Fig. 5F). In cells co-expressing GPERmut and FSHR, 10 nM FSH-treatment induced increases in intracellular cAMP (Fig. 5G), and decreased cell viability when compared to FSH-treated cells co-expressing wild type GPER and FSHR (Fig. 5H). Therefore, inhibition of FSHR-mediated cAMP signaling and increased cell viability is due to a physical interaction between FSHR and GPER (Fig. 5I).

**Fig. 5.**
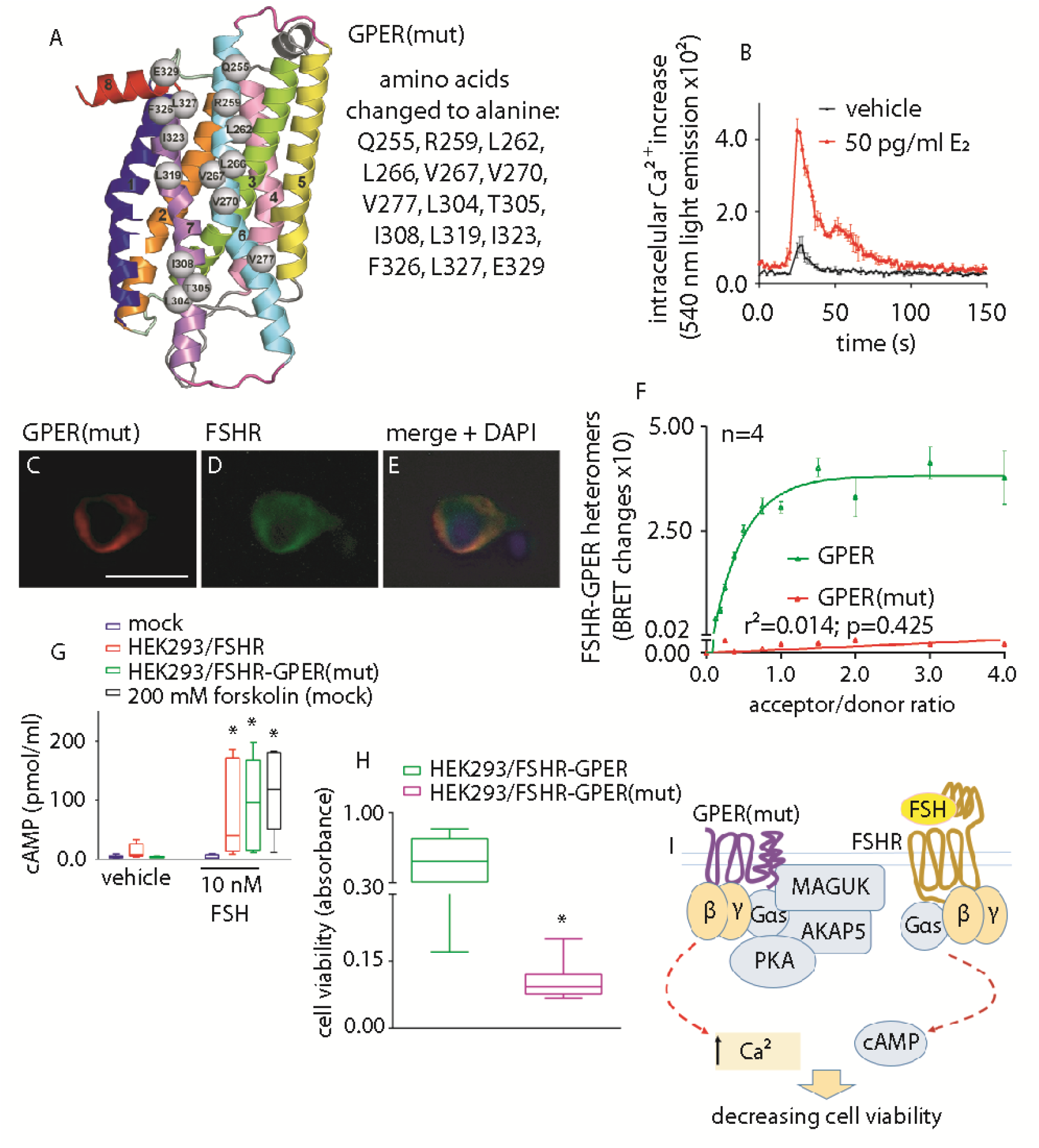
Decrease of cell viability by disruption of FSHR-GPER heteromers. (**A**) Side view, in a direction perpendicular to the bundle main axis, of the GPER1 structural model. The receptor regions are colored as follows: H1, H2, H3, H4, H5, H6, H7, and H8 are, respectively, blue, orange, green, pink, yellow, aquamarine, violet, and red, I1 and E1 are slate, I2 and E_2_ are gray, and I3 and E3 are magenta. The receptor amino acids participating in the interface with FSHR and subjected to alanine replacement are represented as spheres centered on the Cα-atom. They include: Q255, R259, L262, L266, V267, V270, and V277 in H6, L304, T305, I308, L319, I323, F326 and L327 in H7, and E329 in H8. To obtain a GPER(mut) molecule unable to forms heteromers, interacting residues indicated in the box were changed to alanine by *de novo* DNA synthesis. (**B**) Demonstration of GPER(mut) functionality by BRET using the calcium-biosensor, in transiently transfected HEK293 cells. The mutant receptor mediates E_2_-induced intracellular Ca^2+^ increase compared to vehicle, over 150 s (two-way ANOVA; p<0.0001; n=8; means ± SEM). Compounds were injected at the 21-s time-point. (**C-E**) Representative evaluation of GPER(mut) and FSHR co-localization by immunofluorescence, in transfected HEK293 cells. GPER(mut) was detected by the anti-GPER specific primary antibody and TRITC-labelled secondary antibody, while FSHR is identified by the venus tag light emission. Nuclei were blue-stained by DAPI (bar=25 μm). (**F**) FSHR/rluc- and GPER(mut)/venus-tagged proteins do not form heteromers. BRET ratio values resulting from molecular interactions are represented as means ± SEM, together with data from non-mutant GPER (Fig. 2G). Specific association is indicated by data interpolation using linear regression (n=4). (**G**) cAMP increase induced by 10 nM FSH, in HEK293 cells expressing either one or both FSHR and GPER(mut). 50 pg/ml E_2_ were added as indicated, while mock-transfected and forskolin-treated cells served as basal and positive control, respectively. cAMP values are indicated as induced BRET changes over vehicle-treated mocks (*=significantly different *versus* FSH-treated mocks; two-way ANOVA with Tukey’s correction for multiple tests; p≤0.0106; n=8; means ± SEM). (**H**) Comparison of HEK293/FSHR-GPER (Fig. 3I) and HEK293/FSHR-GPER(mut) cell viability, under treatment with 10 nM FSH (*=different *versus* HEK293/FSHR-GPER; Mann-Whitney’s *U* -test; p=0.0003; n=8; means ± SEM). (**I**) Proposed model showing the inability of GPER(mut) to inhibit the activation of FSH-stimulated cAMP production with ensuing decrease of cell viability due to lack of heterodimerisation.

### GPER requires the MAGUK/AKAP5 complex to inhibit FSHR-mediated decrease in cell viability

The role of known GPER-linked machinery^22^ in inhibiting FSHR-mediated cAMP response was evaluated in HEK293 cells via a genome editing approach. *AKAP5* was knocked-out in HEK 293 cells by CRISPR/Cas9 (AKAP5-KO HEK293 cells) (Fig. 6A; data file S9, S10). In these cells, GPER still exhibited heteromer formation with FSHR, as measured by BRET (Fig. 6B). However, FSH treatment of AKAP5-KO HEK293 cells, co-expressing GPER and FSHR, induced intracellular cAMP generation (Fig. 6C), demonstrating AKAP5 is essential in exerting GPER-dependent inhibition of cAMP production occurring via FSHR. Consistent with the cAMP data, the GPER rescue of cell viability requires AKAP5 (Fig. 6D), strengthening the role of this inhibitory machinery in counteracting FSHR activity by GPER. Therefore, one of the molecular mechanisms regulating FSHR-activation of Gαs protein-dependent signals requires the association of GPER and AKAP5, as cells expressing FSHR/GPER but lacking AKAP5 are able to generate cAMP and thus reduce FSH-dependent cell viability (Fig. 6E).

**Fig. 6.**
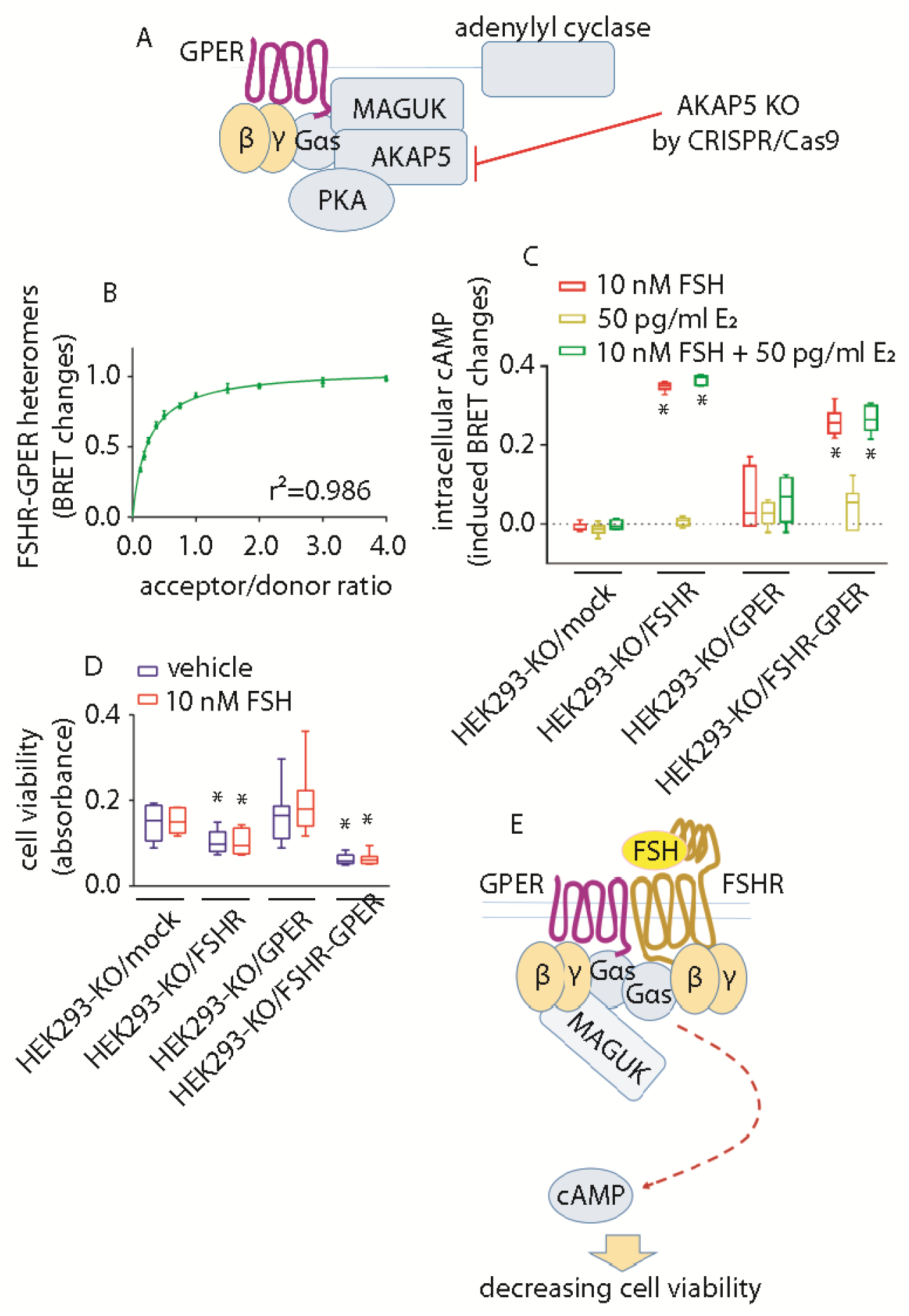
Cell viability decreases upon disrupting the GPER-associated MAGUK/AKAP5 molecular inhibitory complex. (**A**) Representative model showing the development of the AKAP5-KO HEK293 cell line by CRISPR/Cas9. The absence of AKAP5 leads to disruption of the GPER-associated inhibitory machinery without impairing FSHR-GPER heteromers formation. (**B**) FSHR/rluc-GPER/venus-tagged heteromers formation evaluated by BRET, in AKAP5-KO HEK293 cells. Values are expressed as means ± SEM and interpolated by nonlinear regression (n=4). (**C**) Intracellular cAMP increase occurring upon 10 nM FSH-treatment of AKAP5-KO HEK293 cells, transiently expressing FSHR and/or GPER. 50 pg/ml E_2_ were added where indicated and signals acquired by BRET. *=significantly different *versus* FSH-treated mocks; two-way ANOVA with Tukey’s post-hoc test; p<0.0001; n=6; means ± SEM. (**D**) Viability of FSHR- and/or GPER-expressing AKAP5-KO HEK293 cells in the presence and in the absence of 10 nM FSH (*=different *versus* mock; two-way ANOVA with Dunnett’s post-hoc test; p<0.0001; n=8; means ± SEM). (**E**) Model describing the failure of the inhibition of FSHR/Gαs protein signaling and cell viability decrease, in the absence of AKAP5.

### FSH-dependent cell viability and proliferative signals are downregulated in GPER-deficient human granulosa cells: in vivo, pharmacological relevance

The role of GPER in counteracting intracellular death signals delivered through the FSHR was confirmed using human primary granulosa cells, where the effects of FSH were evaluated in the presence and in the absence of GPER via siRNA (Fig. 1C and Fig. 7A-F). Interestingly, GPER-depleted granulosa cells exhibited both increased basal and FSH-induced cAMP production, compared to mock-treated cells (Fig. 7G). Moreover, 10 nM FSH-treatment reduced cell viability following GPER depletion (Fig. 7H) and induced procaspase 3 cleavage as detected by Western blotting (Fig. 7I), suggesting the protective role of GPER from gonadotropin-induced cell death. In these experiments, all samples were maintained under LHCGR depletion via siRNA as LHCGR has been reported to negatively regulate cAMP signaling from FSHR via LHCGR/FSHR heteromers^44^ As increases in intracellular cAMP is known to drive steroid synthesis in granulosa cells, the production of progesterone secreted in FSH-treated cells (Fig. 7J) was also significantly increased in human granulosa cells depleted of GPER (Fig. 7G), thus supporting previously proposed links between steroidogenic and pro-apoptotic pathways in ovarian cells^4,9^.

**Fig. 7.**
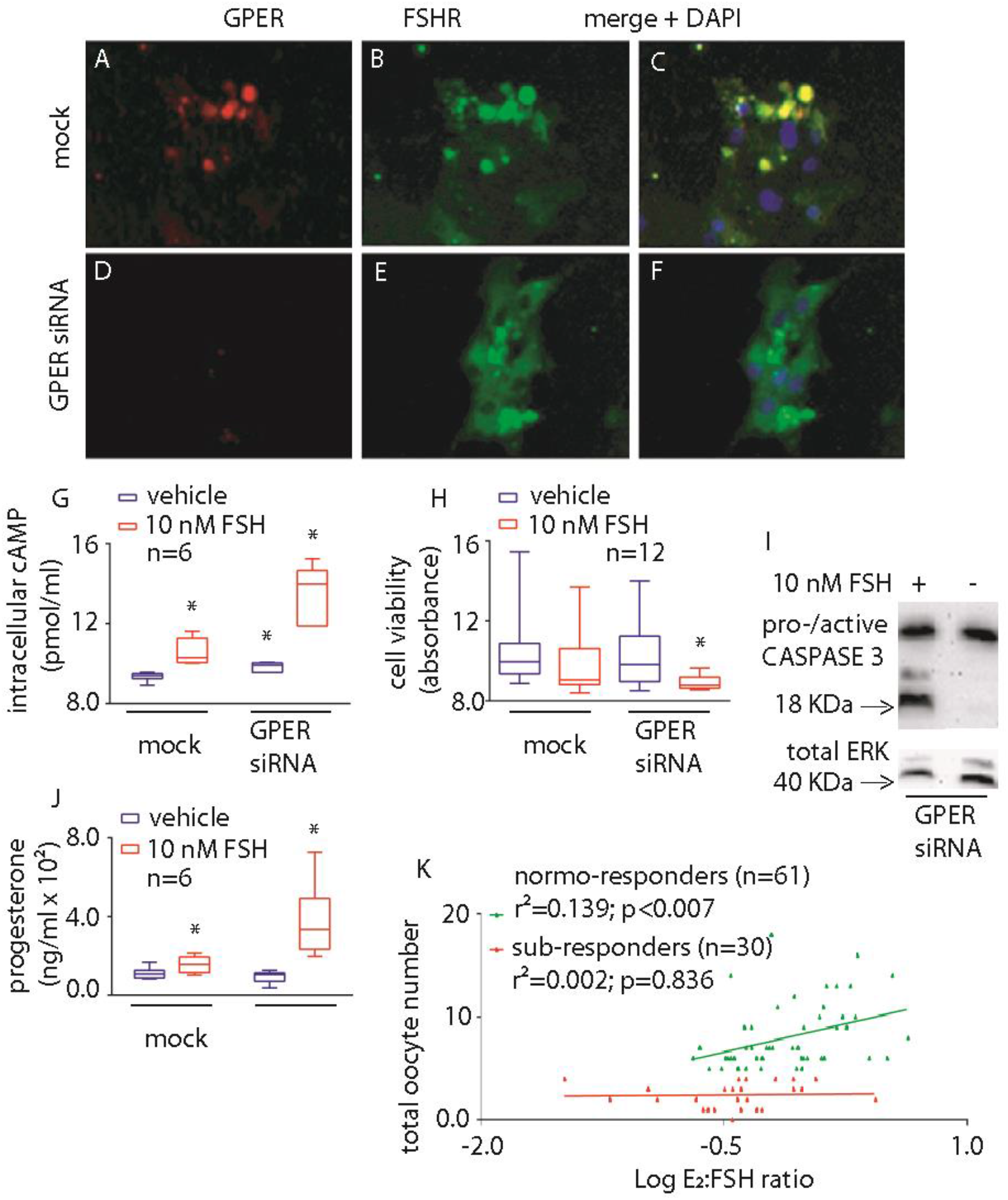
Role of GPER in regulating FSH-mediated cell viability and steroidogenesis in human primary granulosa cells and in patients undergoing FSH stimulation for assisted reproduction. (**A-F**) Representative images of GPER and FSHR co-localization in 48-h mock-(**A-C**) and GPER siRNA-treated (**D-F**) human granulosa cells, detected by immunofluorescence. Specific primary antibody was used for FSHR and GPER binding, as well as TRITC-(**A, D**) and FITC-labelled (**B, E**) secondary antibodies, respectively. Nuclei were blue-stained by DAPI (**C, F**). Bar=25 μm. (G) Intracellular cAMP levels measured in control and GPER siRNA-treated granulosa cells by ELISA, in the presence or absence of 10 nM FSH. Data are represented by box and whiskers plots (*=different *versus* vehicle/mock-treated cells; two-way ANOVA and Sidak’s multiple comparisons test; p≤0.0074; n=8). (H) Granulosa cell viability after 48-h treatment with control/GPER siRNA. Effects of 10 nM FSH were also assessed 24 h before measurements (*=different *versus* vehicle/mock-treated cells; Kruskal-Wallis with Dunn’s correction for multiple tests; p=0.0002; n=12). (I) Evaluation of procaspase 3 cleavage in granulosa cells under 48-h GPER depletion by siRNA. 10 nM FSH was added 24 h before analysis, as indicated, while total ERK was the loading control. (J) Progesterone levels measured in media of control and GPER siRNA-treated granulosa cells, maintained 24 h in the presence or in the absence of 10 nM FSH, by immunoassay. (*=different *versus* vehicle/mock-treated cells; two-way ANOVA and Fisher’s test; p≤0.001; n=6). (K) Correlation between oocyte number and the ratio between E_2_ serum levels and cumulative FSH dose of normo-(n=61) and sub-responder (n=30) women. Patients are represented by points and data were interpolated using linear regression.

This data strongly supports a functional requirement of FSHR-GPER associations in the ovary. As we have shown that proportional amounts of both receptor transcripts correlate with high oocyte yield (Fig. 1D), whereby in FSH sub-responders, low oocyte number is correlated to higher *FSHR* over *GPER* expression, presumably resulting in deleterious effects for cell survival. The concept is further assessed by plotting biochemical parameters against the number of oocytes achieving maturation after controlled ovarian stimulation procedures (Fig. 7K). The ratio between estradiol levels and cumulative FSH dose, assumed to be indicative of the steroid capability to inhibit the selective pressure *via* activation of proliferative and anti-apoptotic signals in growing oocytes, are correlated in normo-responder women while they are not in subresponders who have lower levels of GPER (Fig. 7K). This data further supports our findings in human granulosa cells whereby the reduction of GPER results in enhanced FSH-mediated steroidogenesis but increased loss of cell viability and subsequent decrease in oocyte yield in these patients

## Discussion

This work demonstrates that GPER heteromerization with FSHR shifts the preferential signal transduction from cAMP to pAKT activation, reducing cAMP-dependent apoptosis and favoring cell survival. Our results demonstrate novel aspects of FSHR function, by extending the number of transmembrane partners of FSHR to steroid hormone GPCRs. We also identify the mechanism underpinning GPER-FSHR heteromer inhibition of cAMP signaling is via the GPER-associated MAGUK-AKAP5 protein complex. Critically, our data demonstrate important implications for ovarian physiology and FSH use in fertility treatment, identifying for the first time that FSHR-GPER heteromer represent a druggable target to regulate cell death and survival, and improve fertility outcomes in poor FSH-responders.

The ability of FSHR-GPER to form heteromers was demonstrated complementarily using different approaches and, interestingly, revealed preferential formation of complexes containing a higher number of FSHR than GPER molecules. In that respect, as FSH is thought to bind and activate FSHR trimers^45^, our data could support a role for these asymmetric lower order heterooligomers in regulating cell viability. This inhibitory role of GPER on cAMP is specific to FSHR, since GPER had no effect on LHCGR-mediated cAMP, reflecting structural differences between FSHR and LHCGR, in line with the different and specific physiological role of LH^46,47^ Upon FSHR and GPER co-expression, disruption or structural rearrangements of the FSHR-Gαs protein interaction occurs. These rearrangements are GPER-dependent and negatively impact the FSH-induced cAMP production via the GPER-related anchoring complex *AKAP5*^22^ Data were confirmed in FSHR-GPER co-expressing human primary granulosa cells, where siRNA knockdown of native GPER expression enhanced the FSHR-mediated cAMP steroidogenic pathway^22^ and negatively impacted cell viability. Intact FSHR-GPER complexes responded to FSH treatment by stimulating acute βγ-dependent pAKT activation, which has been associated with cell migration and proliferative events in several cell models^48–51^, and proposed as a target for tumor-suppressing therapies^52^ We found this FSHR-associated pAKT activation occurs relatively rapidly upon cell treatment by FSH, in contrast to what has been previously described^12^ Overall, our findings fulfill the recently proposed criteria for demonstrating functional GPCR heteromers in native tissue: a) receptor co-localization/interaction, b) exhibition of distinct functional properties by heteromers compared to protomers, and c) loss of heteromer-specific properties upon heteromer disruption^53^, in our study this was via knockdown of GPER in primary human granulosa cells; equivalent findings were observed with the GPERmut that specifically could disrupted interactions with FSHR To date, these criteria have only been fulfilled by a small subset of Family A GPCR heteromers^53^.

While proliferative and anti-apoptotic signals are preferentially activated at relatively low FSHR levels^14^, the steroidogenic/pro-apoptotic pathway is stimulated in the presence of increasing receptor number. This is due to concentration-dependent variations of FSHR association affinity with its different intracellular effectors, as shown for other GPCRs^54^, which is similar across different G proteins at low receptor expression levels, while preferential coupling to Gαs protein and activation of cAMP signaling occurs at high receptor expression levels^14^, and now from the findings in this study is likely due to a higher level of FSHR monomers/homomers over FSHR-GPER heteromers. A greater proportion of FSHR only complexes would thus program the cell fate toward FSHR/cAMP-dependent death, a finding well described in the literature^5–7,9,55–59^ but neglected for a long-time due to the lack of evidence on the pro-apoptotic action of FSH *in vivo.* Indeed, FSH action is commonly associated with proliferative events, follicular growth being the physiological example, and as suggested by the presence of FSHR in pathological contexts characterized by uncontrolled cell growth, such as cancer^28^ or endometriosis^29^, which indicated FSHR as a target of anti-cancer drugs^30^.

The proliferative role of FSH is well known, and indeed this hormone is used as a drug for inducing controlled ovarian stimulation and in the clinical setting of infertility treatment^60,61^. On the other hand, the physiological FSH action is directed to estrogen biosynthesis, enhancing cell growth *in vivo*^62^, and it is not surprising that both hormones are placed within proliferative contexts. We suggest that the formation of FSHR-GPER heteromers may be involved in the endocrine regulation of ovarian physiology^63^, providing a molecular mechanism explaining why one follicle becomes dominant although it is exposed to the same hormonal *milieu* of other follicles becoming atretic. Accordingly, FSHR-GPER heteromers would support dominance and further growth, while the lack/insufficiency of GPER expression, and thus GPER-FSHR heteromers, would direct FSH action towards apoptosis in the follicles, which become atretic. Conversely, GPER KO mice feature pathological conditions such as altered glucose and lipid metabolism^64^ and tumorigenesis^65^, without any specific reproductive phenotype^66^. However, the mouse is a multiovulatory species and species-specific mechanisms underlying the endocrine regulation of multi-vs. mono-ovulation are different^67,68^. Whether FSHR-GPER heterodimerization results from the evolution in mono-ovulatory mammals is a topic of future studies.

The ability of these heteromers to modulate opposing apoptotic and proliferative pathways could have implications in hormone-dependent cancers. A number of pro-apoptotic or anti-proliferative actions have been previously associated with FSHR function^3,6^, especially in conditions of high receptor expression levels, similar to other GPCRs^11^. Our data support a mechanism by which FSHR-GPER heteromeric complexes occur and function in certain receptor-expressing tumor cells^27^, inhibiting the pro-apoptotic, cAMP pathway and enhancing the activation of proliferative signals by FSH, thereby upregulating tumor growth. A similar mechanism was previously described for other GPCRs^69,70^. Patients affected by ovarian carcinoma co-expressing FSHR and GPER have lower prognosis than those with cancer cells expressing FSHR or GPER alone^27^ FSHR-GPER heteromerization should be investigated in tumor tissues expressing both receptors, thus representing a potential target for specific drug design. The structural model of the FSHR-GPER interaction predicted, and supported by the ability to disrupt the proposed interface at H6-7, may provide a rationale for future structurebased drug design/discovery.

In terms of clinical applications, our study provides evidence for the potential application of these results in improving outcomes in assisted reproduction and infertility treatment. The possibility to boost follicular growth and maturation in women who are poor responders to ovarian stimulation with FSH (e.g. due to advanced age) is a current challenge in reproductive medicine. Novel biologicals could be designed to specifically and transitorily favor FSHR-GPER oligomerization and, thereby, follicular growth and rescue. Indeed, our data suggest that women who are poor responders to FSH stimulation in an assisted reproduction program, exhibit low expression of GPER that is not correlated with FSHR expression, possibly reflecting reduced ability to form heteromers and thus an insufficient pro-proliferative FSH action in such conditions.

In summary, we have demonstrated that GPER shifts the pro-apoptotic effects of high FSHR expression levels towards upregulation of cell viability. This occurs via formation of heteromeric complexes capable of inhibiting the cAMP pathway but stimulating pAKT, deviating FSHR coupling from Gαs to Gβγ. Our data provides a novel and promising target for improving both infertility treatment and cancer therapy, through the development of drugs favoring or inhibiting FSHR-GPER heteromer function, respectively.

## Materials and Methods

An extended description of materials and methods is available as supplementary material.

### Study design

The objective of this study *in vitro* was to determine the impact of FSHR-GPER heteromers in modulating FSH impact on cell viability. Experiments were performed in biological triplicate, unless otherwise stated, on the basis of previous experiences using transfected cell lines and primary cells *in vitro.* Studies with granulosa cells collected from donor sub- and normo-responder women undergoing oocyte retrieval for assisted reproduction techniques were performed with n=30 and 61 samples, which have a power of about 95% to .detect a difference of 1.7% between the r^2^ values of two groups (alpha=0.05). Experiments were blinded to the operator performing cell handling and real-time PCR analyses. Written consent was collected from women under local Ethics Committee permission (Nr. 796 19th June 2014, Reggio Emilia, Italy). Patients matched these criteria: absence of endocrine abnormalities and viral/bacterial infections, age between 25 and 45 years.

### Statistical analysis

The D’Agostino and Pearson normality test was performed before choosing to use parametric or non-parametric statistics. Two groups of samples were compared using Mann-Whitney’s *U*-test or t-test, while multiple groups were compared using Kruskal-Wallis or one-/two-way ANOVA as proper, as well as proper post-tests and corrections for multiple comparisons depending on the nature of data. Groups were represented in column graph using box and whiskers plots. Non-linear and linear regressions were used for data interpolation in x-y graphs and two linear regressions were compared using the parallelism test of slopes. Statistics were performed using the GraphPad Prism 6.0 software (GraphPad Software Inc., La Jolla, CA, USA).

## Acknowledgments

This study was supported by the Italian Ministry of University and Research (MIUR). Prof. Manuela Simoni, is a LE STUDIUM RESEARCH FELLOW, Loire Valley Institute for Advanced Studies, Orléans & Tours, France, - INRA - Centre Val de Loire, 37380 Nouzilly, France, receiving funding from the European Union’s Horizon 2020 research and innovation programme under the Marie Skłodowska-Curie grant agreement No 665790. We would like to thank Dr Andreas Bruckbauer at the Facility for Imaging of Light Microscopy (FILM), Imperial College London, for technical support with PALM. A.C.H. was supported by grants from the BBSRC (BB/1008004/1) and Genesis Research Trust, N.S.S is supported by an Imperial College London President’s Scholarship.

## Funding

Grant “Departments of Excellence Programme” from MIUR to the Department of Biomedical, Metabolic and Neural Sciences (University of Modena and Reggio Emilia). Polish National Science Centre (NCN) grants: DEC-2015/17/B/NZ1/01777, DEC-2017/25/B/NZ4/02364.

## Author contributions

LC designed the study, managed experiments, performed data analysis and interpretation, wrote the manuscript. CL, EP, SL and LR performed BRET and Western blotting experiments, and data analysis. SS, BM performed BRET and gene expression analysis. SM did immunostainings. CA performed gene expression analysis. NSS have applied the PALM method. JC created the CRISP/Cas9 modified-cells. GB, FP, ALM and MTV provided scientific support, primary cells and tissues, and manuscript editing. ML and GO provided scientific support, data interpretation and manuscript editing. FGK was involved in management of immunostainings and manuscript editing, FF did bioinformatics analyzes, data interpretation and manuscript editing. ARM managed CRISPR/Cas9 experiments, supported data analysis and manuscript editing. ACH supported experiments and study design, provided data interpretation, scientific support, manuscript writing. MS provided study and scientific management, data interpretation and manuscript writing.

## Competing interests

ML and GO are Merck Serono SpA employes without any conflict of interest.

## Notes

### Competing Interest Statement

ML and GO are Merck Serono SpA employes without any conflict of interest. All other authors declare to have no conflict of interests.

## References

1. Correia, S., Cardoso, H. J., Cavaco, J. E. & Socorro, S. Oestrogens as apoptosis regulators in mammalian testis: angels or devils? Expert Rev. Mol. Med. 17, e2 (2015).

2. Lizneva, D. et al. FSH Beyond Fertility. Front. Endocrinol. (Lausanne). 10, 136 (2019).

3. Casarini, L. & Crépieux, P. Molecular Mechanisms of Action of FSH. Front. Endocrinol. (Lausanne). 10, 305 (2019).

4. Amsterdam, A. et al. Alternative pathways of ovarian apoptosis: death for life. Biochem. Pharmacol. 66, 1355–1362 (2003).

5. Amsterdam, A. et al. Steroid Regulation during Apoptosis of Ovarian Follicular Cells. Steroids 63, 314–318 (1998).

6. Casarini, L., Reiter, E. & Simoni, M. ß-arrestins regulate gonadotropin receptor-mediated cell proliferation and apoptosis by controlling different FSHR or LHCGR intracellular signaling in the hGL5 cell line. Mol. Cell. Endocrinol. 437, 11–21 (2016).

7. Aharoni, D., Dantes, A., Oren, M. & Amsterdam, A. cAMP-mediated signals as determinants for apoptosis in primary granulosa cells. Exp. Cell Res. 218, 271–82 (1995).

8. Yoshida, Y. et al. Theophylline and cisplatin synergize in down regulation of BCL-2 induction of apoptosis in human granulosa cells transformed by a mutated p53 (p53 val135) and Ha-ras oncogene. Int. J. Oncol. 17, 227–35 (2000).

9. Breckwoldt, M. et al. Expression of Ad4-BP/cytochrome P450 side chain cleavage enzyme and induction of cell death in long-term cultures of human granulosa cells. Mol. Hum. Reprod. 2, 391–400 (1996).

10. Casarini, L., Santi, D., Simoni, M. & Potì, F. ‘Spare’ Luteinizing Hormone Receptors: Facts and Fiction. Trends Endocrinol. Metab. 29, 208–217 (2018).

11. Revankar, C. M., Vines, C. M., Cimino, D. F. & Prossnitz, E. R. Arrestins block G protein-coupled receptor-mediated apoptosis. J. Biol. Chem. 279, 24578–84 (2004).

12. Gloaguen, P., Crépieux, P., Heitzler, D., Poupon, A. & Reiter, E. Mapping the follicle-stimulating hormone-induced signaling networks. Front. Endocrinol. (Lausanne). 2, 45 (2011).

13. Ulloa-Aguirre, A., Reiter, E. & Crépieux, P. FSH Receptor Signaling: Complexity of Interactions and Signal Diversity. Endocrinology 159, 3020–3035 (2018).

14. Tranchant, T. et al. Preferential β-arrestin signalling at low receptor density revealed by functional characterization of the human FSH receptor A189 V mutation. Mol. Cell. Endocrinol. 331, 109–18 (2011).

15. Gonzalez-Robayna, I. J., Falender, A. E., Ochsner, S., Firestone, G. L. & Richards, J. S. Follicle-Stimulating hormone (FSH) stimulates phosphorylation and activation of protein kinase B (PKB/Akt) and serum and glucocorticoid-lnduced kinase (Sgk): evidence for A kinase-independent signaling by FSH in granulosa cells. Mol. Endocrinol. 14, 1283–300 (2000).

16. Sposini, S. et al. Integration of GPCR Signaling and Sorting from Very Early Endosomes via Opposing APPL1 Mechanisms. Cell Rep. 21, 2855–2867 (2017).

17. Sayers, N. & Hanyaloglu, A. C. Intracellular Follicle-Stimulating Hormone Receptor Trafficking and Signaling. Front. Endocrinol. (Lausanne). 9, 653 (2018).

18. Rossi, V. et al. LH prevents cisplatin-induced apoptosis in oocytes and preserves female fertility in mouse. Cell Death Differ. 24, 72–82 (2017).

19. Chen, J. et al. Gankyrin facilitates follicle-stimulating hormone-driven ovarian cancer cell proliferation through the PI3K/AKT/HIF-1α/cyclin D1 pathway. Oncogene 35, 2506–17 (2016).

20. Revankar, C. M., Cimino, D. F., Sklar, L. A., Arterburn, J. B. & Prossnitz, E. R. A transmembrane intracellular estrogen receptor mediates rapid cell signaling. Science 307, 1625–30 (2005).

21. Heublein, S. et al. The G-protein-coupled estrogen receptor (GPER) is expressed in normal human ovaries and is upregulated in ovarian endometriosis and pelvic inflammatory disease involving the ovary. Reprod. Sci. 19, 1197–204 (2012).

22. Broselid, S. et al. G protein-coupled receptor 30 (GPR30) forms a plasma membrane complex with membrane-associated guanylate kinases (MAGUKs) and protein kinase A-anchoring protein 5 (AKAP5) that constitutively inhibits cAMP production. J. Biol. Chem. 289, 22117–27 (2014).

23. Gonzalez de Valdivia, E., Broselid, S., Kahn, R., Olde, B. & Leeb-Lundberg, L. M. F. G protein-coupled estrogen receptor 1 (GPER1)/GPR30 increases ERK1/2 activity through PDZ motif-dependent and -independent mechanisms. J. Biol. Chem. 292, 9932–9943 (2017).

24. Maggiolini, M. et al. The G protein-coupled receptor GPR30 mediates c-fos up-regulation by 17beta-estradiol and phytoestrogens in breast cancer cells. J. Biol. Chem. 279, 27008–16 (2004).

25. Prossnitz, E. R. & Maggiolini, M. Mechanisms of estrogen signaling and gene expression via GPR30. Mol. Cell. Endocrinol. 308, 32–8 (2009).

26. Pavlik, R. et al. Induction of G protein-coupled estrogen receptor (GPER) and nuclear steroid hormone receptors by gonadotropins in human granulosa cells. Histochem. Cell Biol. 136, 289–99 (2011).

27. Heublein, S. et al. The G-protein coupled estrogen receptor (GPER/GPR30) is a gonadotropin receptor dependent positive prognosticator in ovarian carcinoma patients. PLoS One 8, e71791 (2013).

28. Choi, J.-H., Choi, K.-C., Auersperg, N. & Leung, P. C. K. Overexpression of follicle-stimulating hormone receptor activates oncogenic pathways in preneoplastic ovarian surface epithelial cells. J. Clin. Endocrinol. Metab. 89, 5508–16 (2004).

29. Ponikwicka-Tyszko, D. et al. Functional Expression of FSH Receptor in Endometriotic Lesions. J. Clin. Endocrinol. Metab. 101, 2905–2914 (2016).

30. Perales-Puchalt, A. et al. Engineered DNA Vaccination against Follicle-Stimulating Hormone Receptor Delays Ovarian Cancer Progression in Animal Models. Mol. Ther. 27, 314–325 (2019).

31. Filardo, E. J. A role for G-protein coupled estrogen receptor (GPER) in estrogen-induced carcinogenesis: Dysregulated glandular homeostasis, survival and metastasis. J. Steroid Biochem. Mol. Biol. 176, 38–48 (2018).

32. Barton, M. et al. Twenty years of the G protein-coupled estrogen receptor GPER: Historical and personal perspectives. J. Steroid Biochem. Mol. Biol. 176, 4–15 (2018).

33. Li, Y. et al. FSH stimulates ovarian cancer cell growth by action on growth factor variant receptor. Mol. Cell. Endocrinol. 267, 26–37 (2007).

34. Jeppesen, J. V. et al. LH-receptor gene expression in human granulosa and cumulus cells from antral and preovulatory follicles. J. Clin. Endocrinol. Metab. 97, E1524–31 (2012).

35. Guitart, X. et al. Biased G Protein-Independent Signaling of Dopamine D1-D3 Receptor Heteromers in the Nucleus Accumbens. Mol. Neurobiol. (2019). doi:10.1007/s12035-019-1564-8

36. Ji, I. et al. Trans-activation of mutant follicle-stimulating hormone receptors selectively generates only one of two hormone signals. Mol. Endocrinol. 18, 968–78 (2004).

37. Rivero-Müller, A. et al. Rescue of defective G protein-coupled receptor function in vivo by intermolecular cooperation. Proc. Natl. Acad. Sci. U. S. A. 107, 2319–24 (2010).

38. Jonas, K. C. et al. Temporal reprogramming of calcium signalling via crosstalk of gonadotrophin receptors that associate as functionally asymmetric heteromers. Sci. Rep. 8, 2239 (2018).

39. Tubio, M. R. et al. Expression of a G Protein-coupled Receptor (GPCR) Leads to Attenuation of Signaling by Other GPCRs. J. Biol. Chem. 285, 14990–14998 (2010).

40. Casciari, D., Seeber, M. & Fanelli, F. Quaternary structure predictions of transmembrane proteins starting from the monomer: a docking-based approach. BMC Bioinformatics 7, 340 (2006).

41. Fanelli, F., Seeber, M., Felline, A., Casciari, D. & Raimondi, F. Quaternary structure predictions and structural communication features of GPCR dimers. Prog. Mol. Biol. Transl. Sci. 117, 105–42 (2013).

42. Jonas, K. C., Fanelli, F., Huhtaniemi, I. T. & Hanyaloglu, A. C. Single Molecule Analysis of Functionally Asymmetric G Protein-coupled Receptor (GPCR) Oligomers Reveals Diverse Spatial and Structural Assemblies. J. Biol. Chem. 290, 3875–3892 (2015).

43. Nechamen, C. A., Thomas, R. M. & Dias, J. A. APPL1, APPL2, Akt2 and FOXO1a interact with FSHR in a potential signaling complex. Mol. Cell. Endocrinol. 260-262, 93–9 (2007).

44. Feng, X., Zhang, M., Guan, R. & Segaloff, D. L. Heterodimerization Between the Lutropin and Follitropin Receptors is Associated With an Attenuation of Hormone-Dependent Signaling. Endocrinology 154, 3925–3930 (2013).

45. Jiang, X. et al. Evidence for Follicle-stimulating Hormone Receptor as a Functional Trimer. J. Biol. Chem. 289, 14273–14282 (2014).

46. Casarini, L. et al. Estrogen Modulates Specific Life and Death Signals Induced by LH and hCG in Human Primary Granulosa Cells In Vitro. Int. J. Mol. Sci. 18, 926 (2017).

47. Casarini, L. et al. LH and hCG Action on the Same Receptor Results in Quantitatively and Qualitatively Different Intracellular Signalling. PLoS One 7, e46682 (2012).

48. Ouelaa-Benslama, R. et al. Identification of a GaGßy, AKT and PKCα signalome associated with invasive growth in two genetic models of human breast cancer cell epithelial-to-mesenchymal transition. Int. J. Oncol. 41, 189–200 (2012).

49. Kamal, F. A. et al. Simultaneous adrenal and cardiac g-protein-coupled receptor-gßy inhibition halts heart failure progression. J. Am. Coll. Cardiol. 63, 2549–2557 (2014).

50. Surve, C. R., Lehmann, D. & Smrcka, A. V. A chemical biology approach demonstrates G protein ßy subunits are sufficient to mediate directional neutrophil chemotaxis. J. Biol. Chem. 289, 17791–801 (2014).

51. Matoba, A. et al. The free fatty acid receptor 1 promotes airway smooth muscle cell proliferation through MEK/ERK and PI3K/Akt signaling pathways. Am. J. Physiol. Lung Cell. Mol. Physiol. 314, L333–L348 (2018).

52. Cantley, L. C. & Neel, B. G. New insights into tumor suppression: PTEN suppresses tumor formation by restraining the phosphoinositide 3-kinase/AKT pathway. Proc. Natl. Acad. Sci. U. S. A. 96, 4240–5 (1999).

53. Gomes, I. et al. G Protein-Coupled Receptor Heteromers. Annu. Rev. Pharmacol. Toxicol. 56, 403–25 (2016).

54. Bates, B. et al. Characterization of Gpr101 expression and G-protein coupling selectivity. Brain Res. 1087, 1–14 (2006).

55. Sasson, R., Dantes, A., Tajima, K. & Amsterdam, A. Novel genes modulated by FSH in normal and immortalized FSH-responsive cells: new insights into the mechanism of FSH action. FASEB J. 17, 1256–66 (2003).

56. Sirotkin, A. V et al. Transcription factor p53 can regulate proliferation, apoptosis and secretory activity of luteinizing porcine ovarian granulosa cell cultured with and without ghrelin and FSH. Reproduction 136, 611–8 (2008).

57. Tajima, K. et al. Establishment of FSH-responsive cell lines by transfection of preovulatory human granulosa cells with mutated p53 (p53val135) and Ha-ras genes. Mol. Hum. Reprod. 8, 48–57 (2002).

58. Sirotkin, A. V. et al. cAMP response element-binding protein 1 controls porcine ovarian cell proliferation, apoptosis, and FSH and insulin-like growth factor 1 response. Reprod. Fertil. Dev. 30, 1145–1153 (2018).

59. Maillet, G., Bréard, E., Benhaïm, A., Leymarie, P. & Féral, C. Hormonal regulation of apoptosis in rabbit granulosa cells in vitro: evaluation by flow cytometric detection of plasma membrane phosphatidylserine externalization. Reproduction 123, 243–51 (2002).

60. Santi, D., Potì, F., Simoni, M. & Casarini, L. Pharmacogenetics of G-protein-coupled receptors variants: FSH receptor and infertility treatment. Best Pract. Res. Clin. Endocrinol. Metab. 32, 189–200 (2018).

61. Behre, H. M. Clinical Use of FSH in Male Infertility. Front. Endocrinol. (Lausanne). 10, 322 (2019).

62. Wallach, E. E., Shoham, Z. & Schachter, M. Estrogen biosynthesis—regulation, action, remote effects, and value of monitoring in ovarian stimulation cycles. Fertil. Steril. 65, 687–701 (1996).

63. Hillier, S. G. Current concepts of the roles of follicle stimulating hormone and luteinizing hormone in folliculogenesis. Hum. Reprod. 9, 188–91 (1994).

64. Sharma, G. & Prossnitz, E. R. GPER/GPR30 Knockout Mice: Effects of GPER on Metabolism. Methods Mol. Biol. 1366, 489–502 (2016).

65. Marjon, N. A., Hu, C., Hathaway, H. J. & Prossnitz, E. R. G protein-coupled estrogen receptor regulates mammary tumorigenesis and metastasis. Mol. Cancer Res. 12, 1644–1654 (2014).

66. Prossnitz, E. R. & Hathaway, H. J. What have we learned about GPER function in physiology and disease from knockout mice? J. Steroid Biochem. Mol. Biol. 153, 114–26 (2015).

67. Webb, R. et al. Follicle development and selection: past, present and future. Anim. Reprod. 13, 234–249 (2016).

68. Driancourt, M. A., Webb, R. & Fry, R. C. Does follicular dominance occur in ewes? J. Reprod. Fertil. 93, 63–70 (1991).

69. Moreno, E. et al. Targeting CB2-GPR55 receptor heteromers modulates cancer cell signaling. J. Biol. Chem. 289, 21960–72 (2014).

70. Rozenfeld, R. et al. AT1R-CB_1_R heteromerization reveals a new mechanism for the pathogenic properties of angiotensin II. EMBO J. 30, 2350–63 (2011).

