## supplementary material for "Membrane estrogen receptor (GPER) and follicle-stimulating hormone receptor heteromeric complexes promote human ovarian follicle survival"

**List of supplementary materials**

Supplementary materials and methods

Data file S1

Data file S2

Data file S3

Data file S4

Data file S5

Data file S6

Data file S7

Data file S8

Data file S9

Data file S10

### Supplementary materials and methods

*Cell lines and reagents.*

The HEK293 cell line was available in-house and previously validated for BRET experiments *(71, 72)*. The culture medium was Dulbecco's Modified Eagles Medium (DMEM) enriched by 10% fetal bovine serum (FBS), 2 mM L-glutammine, 100 U/ml penicillin and 50 µg/ml streptomycin. Human primary granulosa cells were collected from ovarian follicles of donor women classified as sub- (≤4 oocytes collected after controlled ovarian stimulation) or normo-responders (>4 oocytes). Cells were handled as previously described *(73)* and cultured in McCoy's 5A medium, 10% FBS, 2 mM L-glutamine, 100 IU/ml penicillin, 100µg/ml streptomycin and 250 ng/ml Fungizone. All culture reagents were from Sigma-Aldrich (Sigma-Aldrich Corporation, St. Louis, MO, USA). Recombinant FSH was provided by Merck KGaA (Gonal-f; Merck KGaA, Darmstadt, Germany) and used at the concentration of 10 nM as previously described *(71)*, while $E_2$ (cat. E8875; 50 pg/ml) *(46)*, 8-br-cAMP (B7880; $1x10^{-15}$-$10^{0}$ nM range) *(71)*, forskolin (F6886; 50 µM) *(6)*, gallein (cat. 371708; 10 µM) *(74)*, fulvestrant (I4409; 2 µg/ml) *(75)* and triton-X (T8787; 5%) were purchased by Sigma-Aldrich. Three human GPER siRNA probes (284631; 3x10 µM/probe) were purchased by ThermoFisher Scientific (Waltham, MA, USA) and delivered into cells using the TransIT TKO transfection reagent (Mirus Bio Corporation, Madison, WI, USA). FSHR-, LHCGR-, $Ca^{2+}$ aequorin- and cAMP CAMYEL-encoding BRET biosensor and cFMS plasmids were available in-house and previously validated *(71–73, 76)*. GPER- and GPER(mut)-encoding BRET biosensor plasmids were developed by *de novo* synthesis and checked by the producer (Gene Universal Inc., Newark, DE, USA), basing on the FLAG/GPER-encoding plasmid *(77)* provided by professor Marcello Maggiolini (University of Calabria, Cosenza, Italy). Gα protein-encoding BRET biosensor plasmids *(78)* were kindly provided by professor Nevin A. Lambert (Augusta University, Augusta, GA, USA). Cell transfections using plasmids were performed using Metafectene PRO (Biontex Laboratories GmbH, Munich, Germany).

*BRET measurements and ELISA.*

Intracellular $Ca^{2+}$ and cAMP levels were evaluated following a validated procedure *(71–73, 79–81)*, in transiently transfected HEK293 cells. G protein coupling experiments were adapted to optimize BRET signals with GPER- and FSHR-encoding plasmids (100 ng plasmid/well as reference amount), starting from the published validation protocol *(78)*. BRET signals were induced using 10 µl/well of 5µM Coelenterazine h (Interchim, Montluçon, France) diluted in 40 µl/well PBS and 1 mM Hepes, in the presence or in the absence of hormones/vehicle. Light emissions were detected at 475±30 and 530±30 nm wavelengths by the CLARIOstar plate reader equipped with a monochromator (BMG Labtech, Ortenberg, Germany). Assessment of protein content was used as loading controls for BRET experiment, while linearity between amount of plasmid administered per well and receptor expressed was confirmed detecting signals of the biosensor-tag. In granulosa cells, cAMP production was measured using an ELISA kit (ab65355; Abcam, Cambridge, UK) following the manufacturer's instructions and signals acquired by a Victor3 plate reader (Perkin Elmer Inc., Waltham, MA, USA).

*Viability assay.*

The 3-(4,5-dimethylthiazol-2-yl)-2,5-diphenyltetrazolium bromide (MTT) assay was performed according the manufacturer's instruction and previous optimizations *(6, 46)*. The assay solution was prepared starting from the powder commercially available (M5655; Sigma-Aldrich) and left into cell well-plates 4 h before to be lysed with isopropanol and determine absorbance value by a plate reader.

*PD-PALM imaging and localization analysis*.

PD-PALM imaging was carried out as previously described *(42)*. Briefly, anti-FLAG and anti-HA antibodies were labelled with CAGE 500 and 552 photoswitchable dyes, respectively, following manufacturer's protocol (Abberior GmbH, Göttingen, Germany). Degree of labelling was determined to be 1.0 ± 0.2 dye molecules per antibody and 1.3 ± 0.1 dye molecules per antibody for for FLAG-CAGE 500 and HA-CAGE 552 respectively. HEK293 cells were transfected with 0.5 µg FLAG-tagged GPER and 1 µg HA-tagged FSHR per well of a 6-well plate. Transfected cells were seeded onto 1.5 glass bottom dishes (MatTek Corporation, Ashland, MA, USA). Cells were incubated with CAGE-conjugated antibodies at 37 °C for 30 min. Cells were washed with PBS and fixed in 4% paraformaldehyde with 0.2% glutaraldehyde for 30 min. Cells were washed with DPBS and maintained in DPBS in the dark until imaging. Single color or simultaneous dual-color images were acquired using an Elyra PS1 (Carl Zeiss AG, Oberkochen, Germany). Images were obtained at 100x oil emersion, 1.45 NA objective. Photo-conversion of CAGE 500 and 552 dyes was achieved with 405 nm light source and was simultaneously imaged and photo-bleached by 491 and 561 nm lasers, respectively. Acquisition of images is as previously described *(42)*.

Analysis was carried out on cropped non-overlapping 7 × 7 µm regions, within cell-cell boundaries, from 491- and 561-nm channels and analyzed for localized receptors by QuickPALM Fiji plugin to generate x-y coordinates of localized receptors in each channel. The number of associated protomers derived from the *x-y* coordinates were quantified using a custom Java application (PD-Interpreter) *(42)*. The percentage and protomer composition of FSH/GPER homomers and heteromers was carried out using a second order Getis Franklin neighbourhood analysis with a search radius of 50 nm. Outputted data was represented as a co-localization plot with heat maps generated to represent the different numbers of associated protomers observed.

*Protein analysis*.

Protein content of samples was determined by the colorimetric Bradford assay using a commercial reagent and following the manufacturer's instructions (#5000201; Bio-Rad Laboratories Inc., Hercules, CA, USA), then 595-nm signals were acquired by a plate reader. Target proteins were analyzed by 12% SDS-PAGE and Western blotting using a validated protocol *(6, 46, 47, 54)* after extraction in ice-cold RIPA buffer along with PhosStop phosphatase inhibitor and a protease inhibitor cocktail (Roche, Basel, Switzerland). Human pAKT, -pCREB and total ERK (#9271, #9198, #4695, respectively; Cell Signaling Technology Inc., Danvers, MA, USA), active/pro caspase 3 (#MA1-91637; Thermo Fisher Scientific, MA, USA), FSHR (#PA5-28764; ThermoFisher Scientific, Waltham) and GPER (#AF5534; Bio-Techne Corporation, Minneapolis, MN, USA) were evaluated using specific antibodies and secondary anti-rabbit (#NA9340V; GE HealthCare) or -goat HRP-conjugated antibodies (#ab6885; Abcam), as appropriate. A mouse HRP-conjugated anti-human β-ACTIN antibody (#A3854; Sigma-Aldrich) was also used. Signals were developed with ECL (GE HealthCare) and detected VersaDoc system using the QuantityOne analysis software (Bio-Rad Laboratories Inc.).

*Steroid hormone measurements.*

Serum FSH levels were calculated as the cumulative dose injected in patients throughout the ovarian stimulation period. Total progesterone and estradiol was measured in sera and in the cell media ($4\times10^4$ cells/well), as indicated, by an immunoassay analyzer (ARCHITECT second Generation system; Abbot Diagnostics, Chicago, IL, USA) after freezing-thawing samples *(73)* and data normalized over cell amount.

*Structural modeling.*

No crystallographic structures are available so far for GPER. Structural models of the two receptors (human species) deprived of the N-terminal and C-terminal regions were achieved by comparative modeling (by the Modeler software *(82)*) by using the crystal structure of an inactive state of the μ-opioid receptor (PDB: 4dkl) as a template, according to a protocol already described *(83)*. As for the FSHR, to model the insertions, the following portions: 488-490 (in E1), 580-583 (in E2), 617-621 and 628-631 (both in I3), and 669-671 (in E3) were deleted in the template crystal structure, followed by addition of external α-helical restraints to the following amino acid stretches: 421-432 (extracellular extension of H2), and 552-559 (cytosolic extension of H6). As for GPER to model the insertions, the following portions: 219-230 (C-terminal portion of E2 and initial portion of H5), 222-225 (in E2), 259-263 and 270-273 (both in I3), and 307-311 (in E3) were deleted in the template crystal structure, followed by addition of external α-helical restraints to the following amino acid stretches: 52-62 (cytosolic half of H1), 226-233 (cytosolic extension of H5), 241-244 (cytosolic extension of H6), 271-278 (extracellular extension of H6), and 286-293 (extracellular extension of H7). For each receptor, one-hundred models were built by randomizing all the Cartesian coordinates of standard residues in the initial model. The best model according to quality checks was subjected to application of rotamer libraries to those side chains in non-allowed conformation.

Prediction of likely architectures of FSHR-GPER heterodimer followed a computational approach developed for quaternary structure predictions of transmembrane α-helical proteins, defined as a FiPD-based approach *(40, 41)*. It consists in rigid-body docking using a version of the ZDOCK program devoid of desolvation as a component of the docking score (v2.1) *(84)*. FSHR was used as a fixed protein (target) and GPER as a mobile protein (probe) and vice versa in two distinct docking runs. A rotational sampling interval of 6º was set (i.e., dense sampling) and the best 4000 solutions were retained and ranked according to the ZDOCK score. Such solutions were filtered according to the "membrane topology" filter (by using the FiPD software *(40)*, which discards all those solutions that violate the membrane topology requirements. The membrane topology filter, indeed, discards all the solutions characterized by a deviation angle from the original z-axis, i.e. tilt angle, and a displacement of the geometrical centre along the z-axis, i.e. z-offset, above defined threshold values, which were 0.4 radians and 6.0 Å, respectively. The filtered solutions from each run were merged with the target protein, leading to an equivalent number of dimers that were clustered using a $C_\alpha$-RMSD threshold of 3.0 Å for each pair of superimposed dimers. All the amino acid residues in the dimer were included in $C_\alpha$-RMSD calculations. Cluster analysis was based on a QT-like clustering algorithm *(85)* implemented both in the FiPD and Wordom software *(40, 86)*. Since the filtering cutoffs of the membrane topology parameters are intentionally quite permissive, inspection of the cluster centres (i.e. the solutions with the highest number of neighbours in each cluster) served as a final filter to discard remaining false positives, thereby leading to a dramatic reduction of the reliable solutions. The best scored docking solutions from the most populated and reliable clusters were finally considered. Cluster reliability was based on the MemTop score, accounting for the goodness of the membrane topology. Such index is defined according to the following formula:

$$MemTop = \sqrt{\langle tilt_{nor} \rangle^2 + \langle Z_{offnor} \rangle^2}$$

where, the squared terms are, respectively, the normalized tilt angle and the z-offset averaged over all the members of a given cluster. Normalization of each tilt angle and z-offset value was carried out by dividing each value for the respective cutoff value, i.e. 0.4 radians, for the tilt angle, and 6.0 Å, for the z-offset. The optimal value for such index is zero.

Final selection of the likely heterodimer relied on a consensus from the two different docking runs.

*CRISPR/Cas9 experiments*.

Since AKAP5 is encoded by a single exon (exon 2), two guide RNA probes (gRNAs) were used to excise the entire genetic region. gRNAs were designed using Benchling platform and then ordered as a synthetic DNA fragment (Gene Strings, Thermo Fisher Scientific) flanked by two restriction enzyme BbsI recognition sites. Each of the two double-stranded DNA fragment was then cloned into a pSpCas9(BB)-2A-Puro (PX459) V2.0 vector, respectively, which was a gift from Feng Zhang (#62988; Addgene, Watertown, MA, USA), using digestion/ligation protocol *(87)*. The vectors were then amplified and purified from ampicillin-resistant bacteria with Zyppy Mini Plasmid Kit (Zymo Research, Irvine, CA, USA).

HEK293 cells transfection was performed using TurboFect reagent (Thermo Fisher Scientific) according to manufacturer protocol. After 48 h, we enriched cell media by 1.2 µg/ml puromycin before to verify the KO by PCR reaction in single clone lysates.

*Immunofluorescence and immunohistochemistry*.

FSHR and GPER expression was detected by immunofluorescent microscopy in both granulosa and 48-h transfected HEK293 cells using a ZOE Fluorescent Cell Imager (Bio-Rad Laboratories Inc.). Granulosa cells were fixed by 3-min treatment with 4% ice-cold paraformaldehyde/PBS, heated 20 s x 600 W in a microwave and incubated with 20 µg/ml rabbit anti-FSHR (#PA5-28764; ThermoFisher Scientific) and/or 20 µg/ml goat anti-GPER (#ab6885; Abcam) primary antibodies. Secondary antibodies were donkey anti-rabbit FITC and anti-goat TRITC (#ab6798 and #ab6882; Abcam). In transfected HEK293 cells, treatment by the anti-GPER and proper secondary antibody was performed, while FSHR was identified by the venus-tag emission. 300 nM DAPI was used for staining nuclei.

Immunohistochemistry was performed using an R.T.U. Vectastain Universal Elite ABC kit and DBA Substrate Kit (both from Vector Laboratories, Burlingame, CA, USA) according to the manufacturer's instructions. Briefly, paraffin-embedded ovarian tissues of fertile women at the follicular stage, stored in a pathological anatomy laboratory, were sliced into 5 µm sections, deparaffinized and hydrated using decreasing concentration of ethanol. Antigen retrieval was performed by heating sections for 10 min in sodium citrate buffer (pH = 6.0), followed by cooling at room temperature. After treatment for 10 min with 3% hydrogen peroxide in deionized water to quench endogenous peroxidases and for 30 min with normal horse serum (NHS) (Vector Laboratories, Burlingame, CA), sections were incubated overnight at 4°C with primary antibodies diluted in 0.3% NHS in PBS: 20 µg/ml rabbit anti-FSHR (#PA5-28764;

ThermoFisher Scientific, Waltham, MA, USA) or 20 µg/ml rabbit anti-GPER (#LS-A4271; LifeSpan BioSciences Inc., Seattle, WA, USA). Control sections were incubated with 0.3% NHS in PBS in the absence of primary antibody. Slides were incubated in biotinylated horse anti-rabbit (Vector Laboratories, Burlingame, CA) for 30 min at RT followed by 30 min in Elite ABC reagent (Vector Laboratories, Burlingame, CA). Visualization of antibody binding was achieved applying DAB solution (Vector Laboratories, Burlingame, CA) for 5 min. Sections were then counterstained with hematoxylin, dehydrated and mounted using a non-aqueous mounting medium. Images were acquired by Zeiss Axioplan 2 microscope equipped with a Nikon Digital Sight camera.

*Gene expression and DNA sequencing analysis*.

Gene expressions were evaluated using specific primer sequences. *FSHR* (NM_000145.3): fwd 5′-GGAGGTGATAGAGGCAGATG-3′; rev 5′-GGGTTGATGTAGAGCAGGT- 3′; *GPER* (AF027956): fwd 5’-CTGAACCGCTTCTGTCAC-3’; rev 5’-ACTGCTGAACCTCACATC-3’. The *RPS7* housekeeping gene was the loading control (NM_001011.4; fwd: 5'-AATCTTTGTTCCCGTTCCTCA-3'; rev 5'-CGAGTTGGCTTAGGCAGAA-3'). Primers were validated by PCR and DNA Sanger sequencing (data file S2), performed using known settings *(71)*.

*Flow Cytometry*.

HEK293 cells were transiently transfected with either plasmid encoding for empty pcDNA3.1 vector (mock) or increasing concentrations of FLAG-tagged FSHR- or untagged GPER-encoding plasmids. To evaluate the relative expression and localization of FSHR and GPER proteins in the cell membrane, cells were also co-transfected with a fixed concentration of FLAG-tagged FSHR- and increasing concentrations of GPER-encoding plasmids. Cells were detached, washed and resuspended in working buffer (PBS without $Ca^{2+}$ and $Mg^{2+}$; 1% BSA, 2 mM EDTA) 48 hours after transfection. Then, cells were incubated with phycoerythrin (PE)-conjugated anti-FLAG antibody (anti-DYKDDDDK-PE, #130-101-577; 1:100 dilution; Milteny Biotech, Bergisch Gladbach, Germany), 1 h at 4°C, to reveal membrane expression of FSHR. GPER cell membrane expression was revealed by incubating transfected cells 2 h at 4°C with anti-GPER primary antibody (#LS-A4271; LifeSpan BioSciences Inc.) followed by incubation with secondary antibody Alexa Fluor 647-conjugated AffiniPure (1:100 dilution; #AS075 Jackson Laboratory, Bar Harbor, USA), 1 h at 4°C. Cells were washed twice and re-suspended in working buffer before analysis with MACSQuant Analyzer 10 Flow cytometer (Milteny Biotech). Data were analysed and plotted with the FlowJo software (FlowJo, Ashland, OR, USA).

**Supplementary Materials**

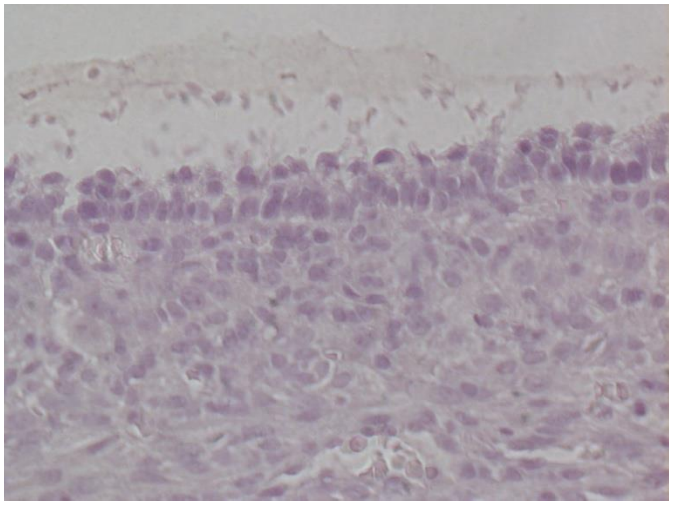

Data file S1. Control section of IHC for FSHR and GPER. The same area was analyzed in serial sections, respectively incubated with anti-FSHR or anti-GPER antibodies or in absence of primary antibody (control). No staining was observed in this area indicating the specific signal of FSHR and GPER.

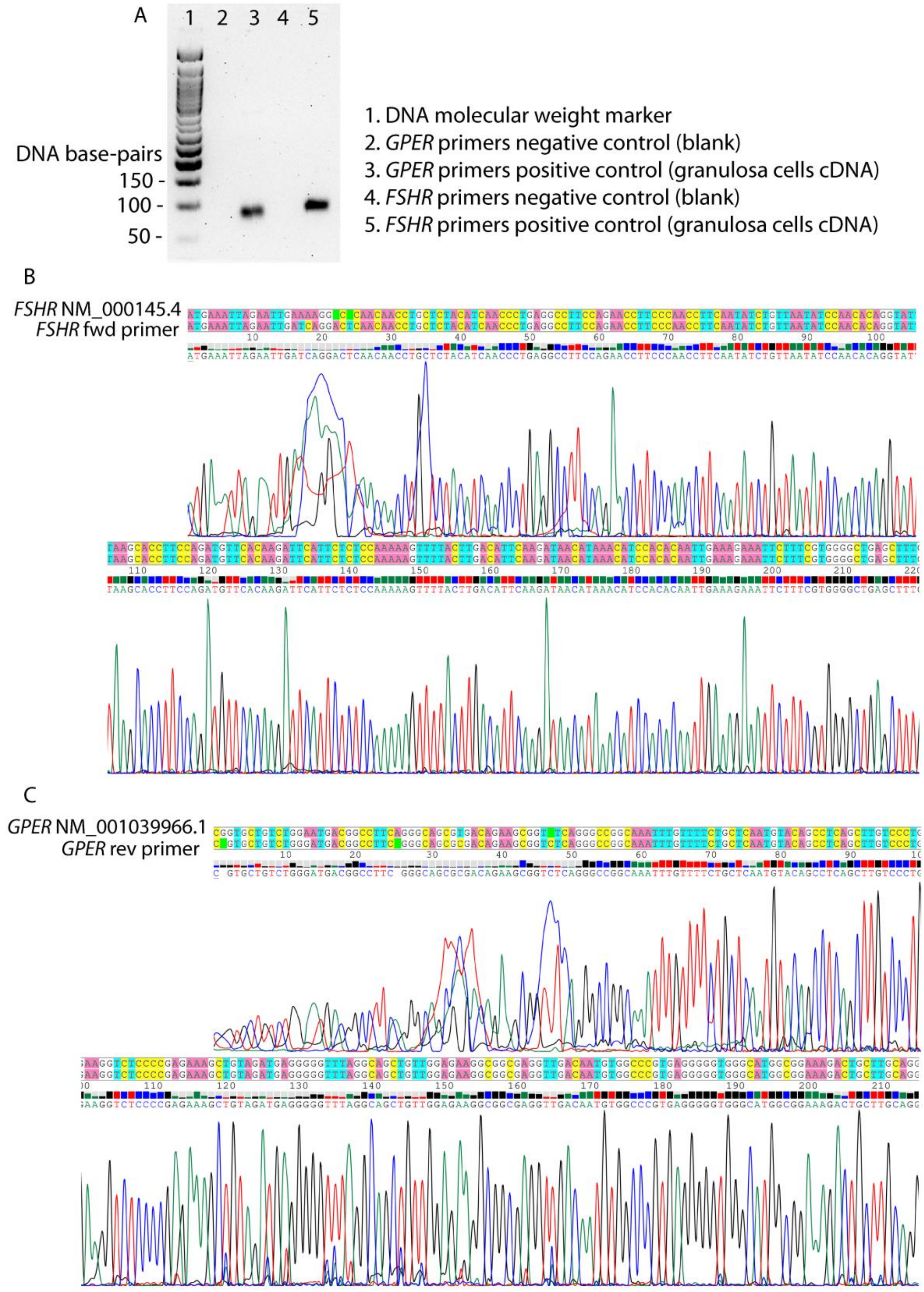


Data file S2. Validation of the *FSHR* and *GPER* primer sequences used for real-time PCR analyses. (**A**) Control experiment performed using blank samples and cDNAs from human primary granulosa cells. Analysis by PCR and 1% agarose-gel electrophoresis demonstrates the presence of bands at the predicted molecular weights. (**B**, **C**) Images of partial electropherograms obtained by DNA Sanger's sequencing demonstrating the specificity of the primers used for real-time PCRs displayed in the figure 6 (only one of the two primers per gene is shown). FSHR and GPER BRET plasmids were used as templates.

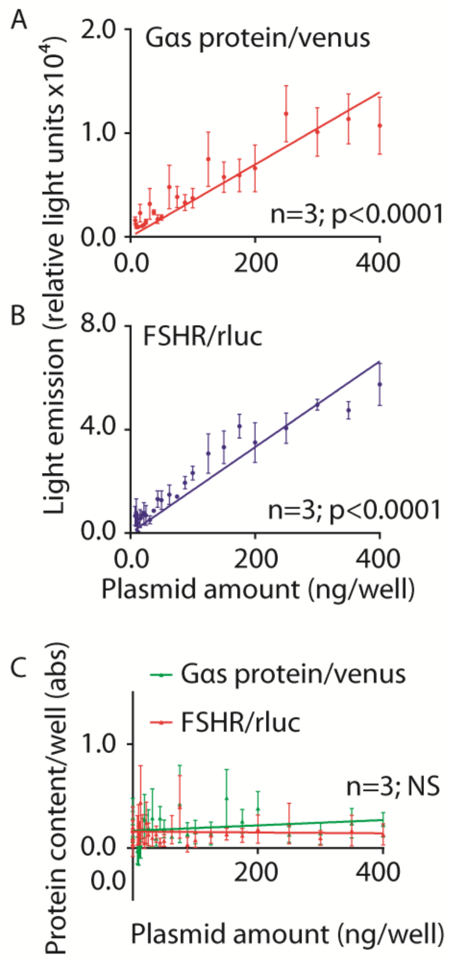


Data file S3. Control of cell transfection efficiency using FSHR and Gαs BRET biosensors. (**A**, **B**) Linear correlation between amount of Gαs protein/venus- or FSHR/rluc-encoding plasmid administered per well, and amount of protein by transiently transfected HEK293 cells. Light emitted by the biosensors was measured by BRET and coelenterazine H was added as a substrate in samples expressing the FSHR/rluc-encoding plasmid 5 min before signal acquisition. Data (means ± SEM; n=3) were interpolated by linear regression forced to pass through x=0.0 and y=0.0. (**C**) Bradford's assay of HEK293 cells transfected with increasing concentrations of Gαs protein/venus- or FSHR/rluc-encoding plasmid (BRET loading control). Protein content was determined by 595 nm absorbance (means ± SEM; n=3) and data interpolated by linear regression.

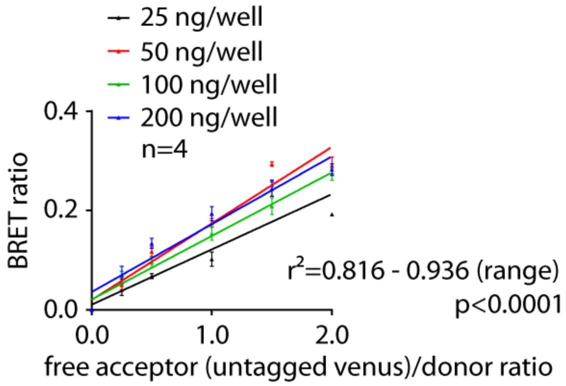


Data file S4. Negative control of FSHR/rluc and venus BRET signal specificity. HEK293 cells were transfected with the indicated concentrations of FSHR/rluc- and with increasing amount of untagged venus-encoding plasmid, then BRET signals were acquired by a plate reader and plotted as means ± SEM against acceptor/donor ratio (n=4). Data were interpolated by linear regression demonstrating the unspecific interaction between FSHR/rluc and untagged venus molecules.

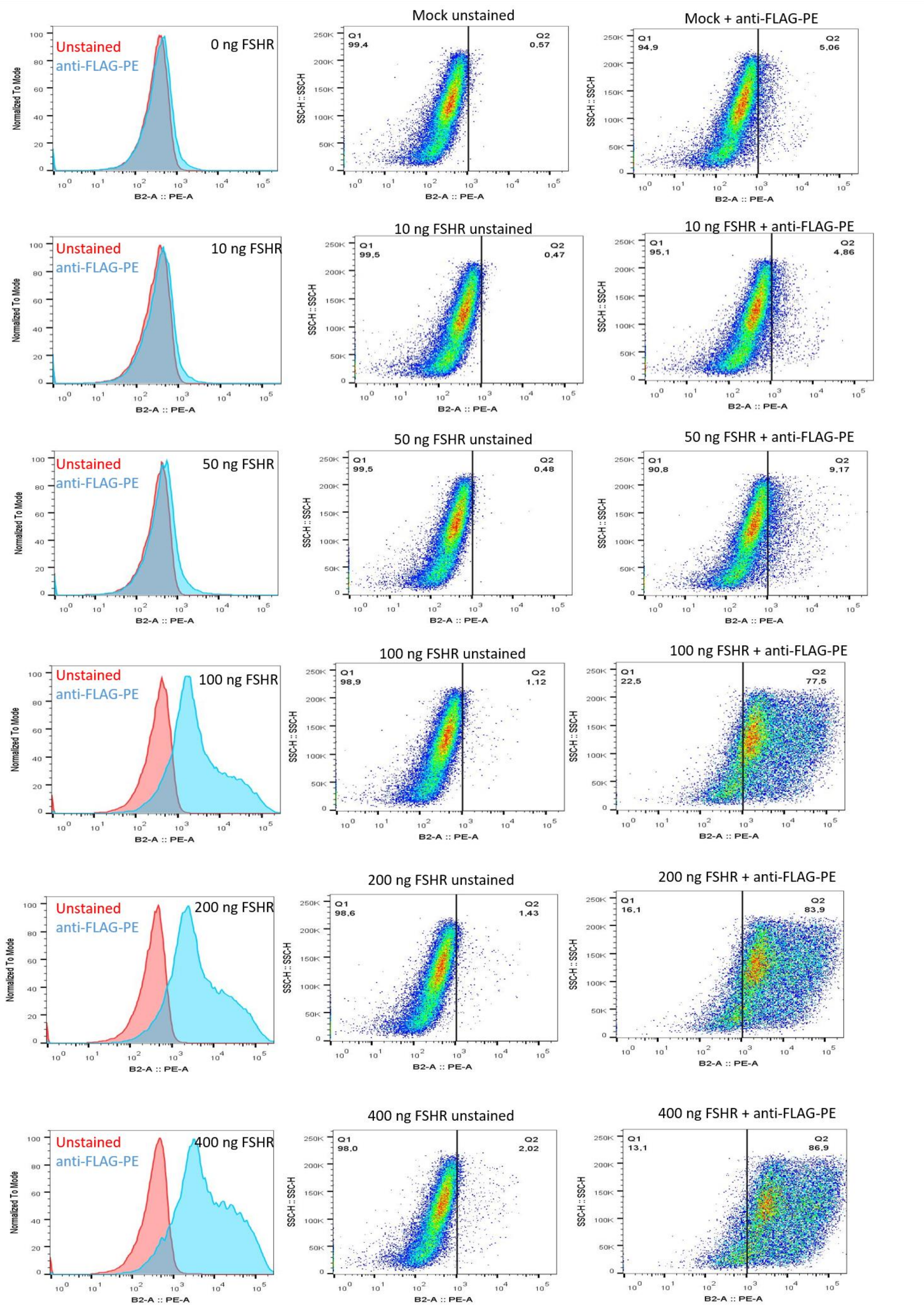


Data file S5 Flow cytometry analysis of plasma membrane FSHR expression levels at different concentrations. 5 x $10^5$ HEK293 cells were transfected either with mock vector or increasing concentrations of FLAG-FSHR-encoding plasmid (10-400 ng/well). Then, cells were stained with anti-FLAG-PE antibody for detection of FSHR and analyzed by flow cytometry. Red peaks in histograms refer to unstained cells while light blue peaks refer to cells incubated with anti-FLAG-PE. Total number of cells in each peak was normalized to 100 % (normalized to mode). Dot-plots show side-scatter versus PE-intensity. Q1 represents the percentage of unstained cells and Q2 the percentage of stained cells for each dot-plot.

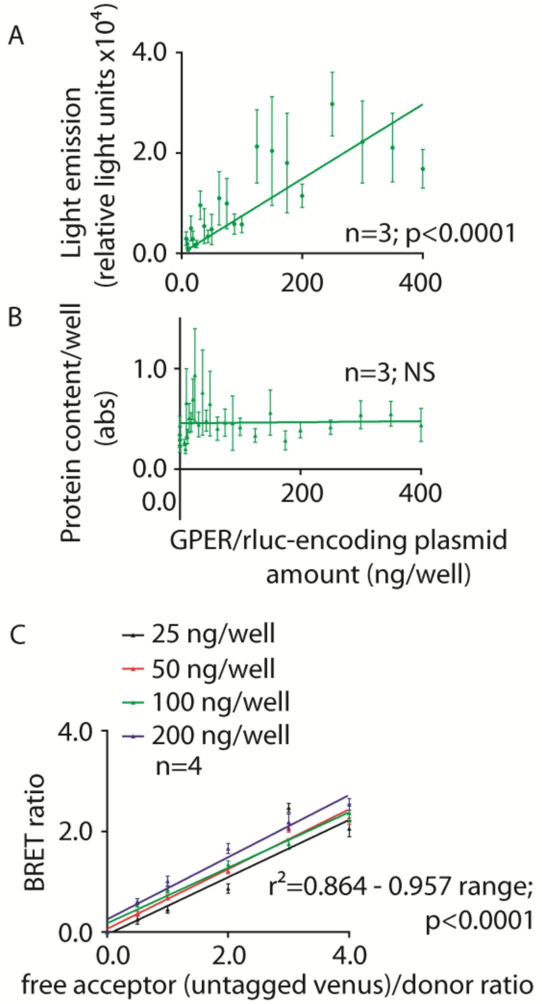


Data file S6. Control of cell transfection efficiency and signal specificity using the GPER BRET biosensors. (**A**) Linear correlation between amount of GPER/rluc-encoding plasmid per well and protein encoded, in transfected HEK293 cells. Light emitted by the biosensors was measured by BRET 5 min after addition of coelenterazine H. Data were interpolated by linear regression forced to pass through x=0.0 and y=0.0 (means ± SEM; n=3). (**B**) BRET loading control determined by Bradford's assay. HEK293 cells were transfected with increasing concentrations of GPER/rluc-encoding plasmid and the protein content was detected (absorbance at 595 nm), plotted as means ± SEM against the amount of plasmid per well and interpolated by linear regression (n=3). (**C**) Negative control of GPER/rluc and venus BRET signal specificity. HEK293 cells were transfected with the indicated concentrations of GPER/rluc- and with increasing amount of untagged venus-encoding plasmid, then BRET signals were acquired and plotted against the acceptor/donor ratio (means ± SEM; n=4). Data interpolation by linear regression demonstrates the unspecific interaction between GPER/rluc and untagged venus molecules.

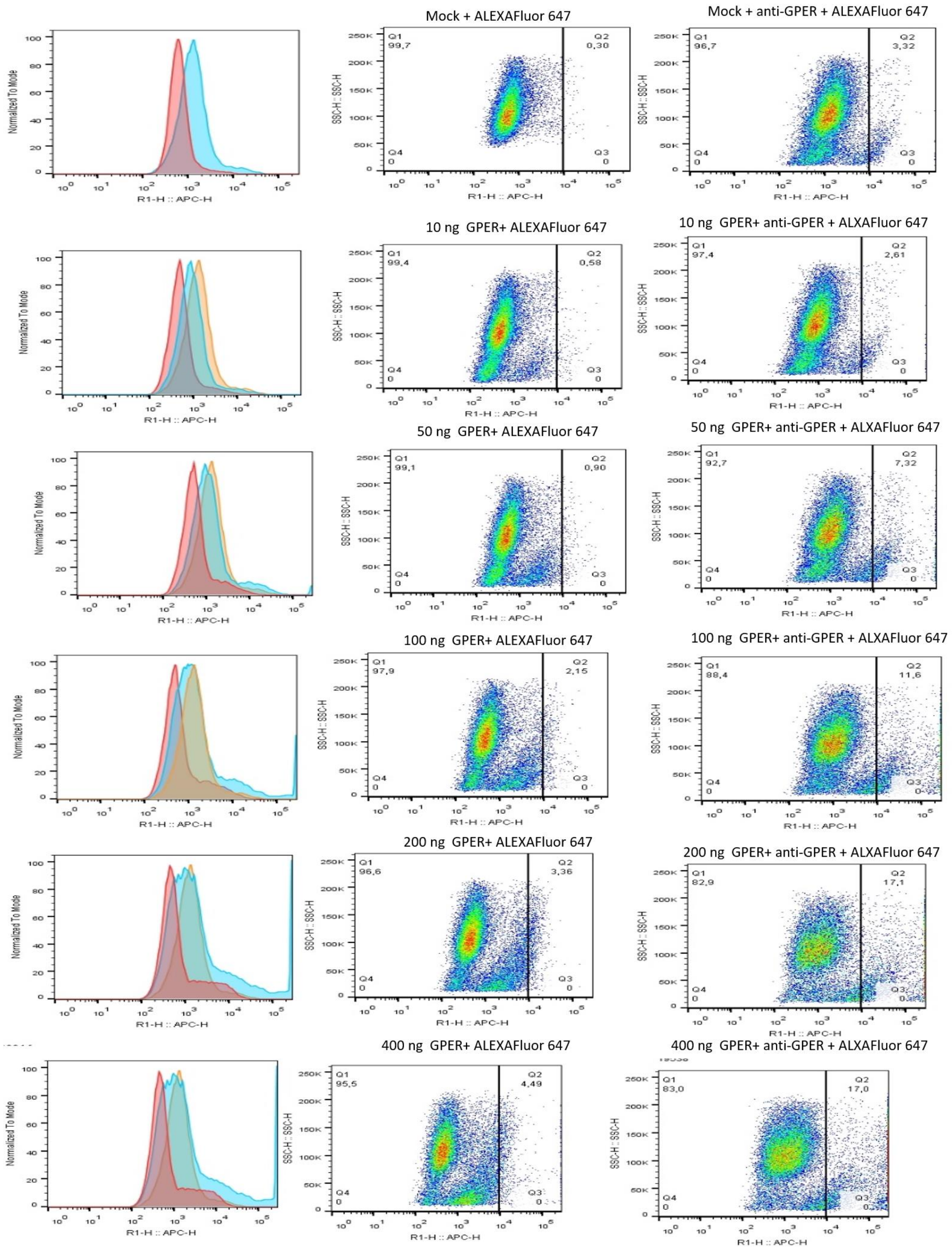


Data file S7. Flow cytometry analysis of plasma membrane GPER expression levels at different concentrations. 5 x $10^5$ HEK293 cells were transfected either with mock vector or increasing concentrations of GPER-encoding plasmid (10-400 ng). Then, cells were incubated with anti-GPER primary antibody followed by incubation with ALEXA Fluor 647 secondary antibody and analyzed by flow cytometry. Red peaks in histograms refer to cells incubated with secondary antibody only, light blue peaks refer to cells incubated with primary and secondary antibody while orange peaks refer to mock-transfected cells incubated with primary and secondary antibodies. Total number of cells in each peak was normalized to 100 % (normalized to mode). Dot-plots show side-scatter versus ALEXA Fluor 647 (APC-H) intensity. Q1 represents the percentage of cells negative to ALEXA FLUOR 647 staining and Q2 the percentage of cells positive to ALEXA FLUOR 647 staining in each dot-plot.

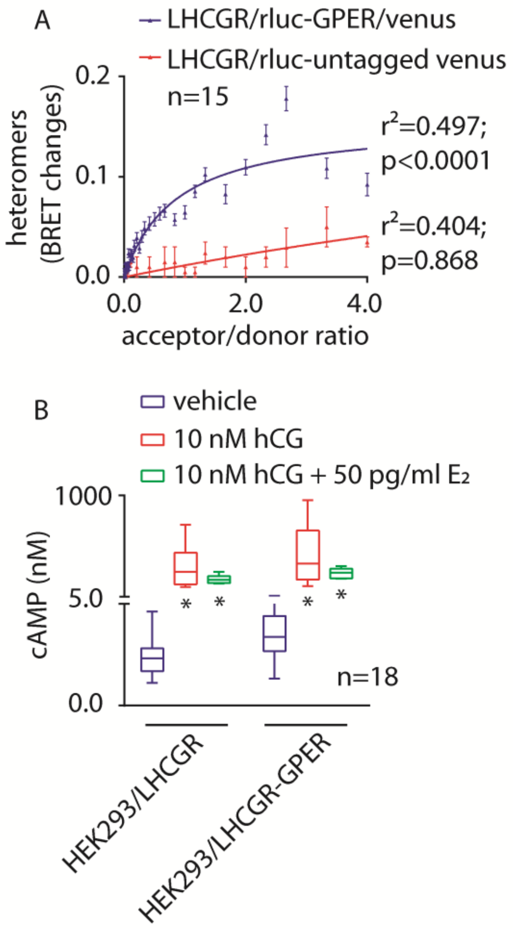


Data file S8. hCG-induced cAMP increase in the presence of LHCGR and GPER heteromers. (**A**) BRET signal demonstrating the formation of LHCGR/rluc- and GPER/venus-tagged heteromers, in transfected HEK293 cells. BRET ratio values resulting from molecular interactions were represented in the x-y graph as means ± SEM (n=5). Specific binding is indicated by data interpolation using non-linear regression, which results in the logarithmic curve, while unspecific binding between LHCGR/rluc and untagged venus molecules is indicated by linear regression. (**B**) 10 nM hCG-induced intracellular cAMP increase, in HEK293 cells expressing either one or both LHCGR and GPER. 50 pg/ml $E_2$ was added as indicated. cAMP was measured by ELISA and represented by box and whiskers plots (*=significantly different *versus* vehicle-treated HEK293/LHCGR; two-way ANOVA with Sidak's correction for multiple tests; p<0.0001; n=18).

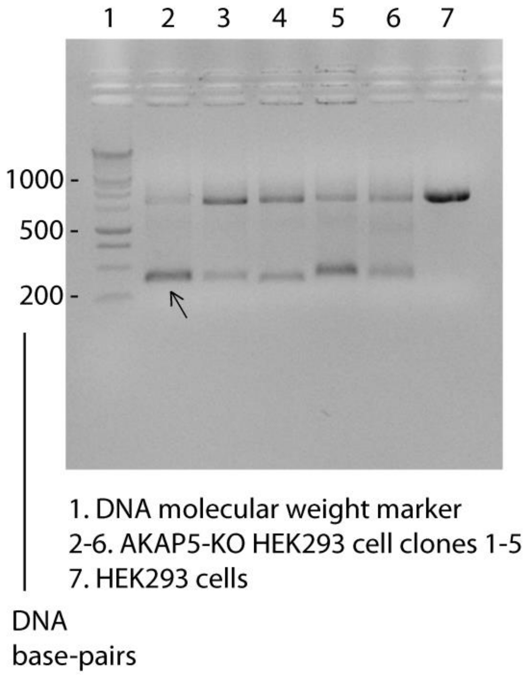


Data file S9. Screening of *AKAP5*-KO HEK293 cells by PCR and 1% agarose-gel electrophoresis. *AKAP5* forward (fwd) and reverse (rev) primers (*GGAGTAAGATGAAAGGTATGAATATGCC* and *CTGCAATCTGTGCTGACTTCC*, respectively; 58°C melting temperature) were designed using the human gene sequence as a template (NC_000014.9). PCR reactions were performed using genomic DNAs extracted from five KO clones (lanes 2-6), while DNA from "native" HEK293 cells was used as a control (lane 7). The predicted sequenced amplified in *AKAP5*-KO cells is of 335 base-pairs, while a band of about 900 base-pairs is predicted to be amplified in the WT *AKAP5*-positive sample. A variable grade of *AKAP5*-WT cell contamination persists among the cultured *AKAP5*-KO HEK293 cells maintained under selective pressure by 1.2 µg/ml puromycin. Arrow indicates the clone used for experiments (Fig. 5).

Data file S10. AKAP5 gene and CRISPR/Cas9 targeting. The target gRNAs are depicted in red. The entire coding region in exon 2 was erased. Sequence of the Cas9 backbone carried in pSpCas9(BB)-2A-Puro (PX459) V2.0 vectors are below and gRNA sequences indicated in bold.

5'-
AMMTCGMMAWWACGATACAAGCTGTTAGAGAGATAATTGGAATTAATTTGACTGT
AAACACAAAGATATTASTACAAAATACGTGACGTAGAAAGTAATAATTTCTTGGGTA
GTTTGCAGTTTTAAAATTATGTTTTAAAATGGACTATCATATGCTTACCGTAACTTGA
AAGTATTTCGATTTCTTGGCTTTATATATCTTGTGGAAAGGACGAAA**CACCGATCAG**
**CAGAAGGTAGTCCTG**GTTTTAGAGCTAGAAATAGCAAGTTAAAATAAGGCTAGTC
CGTTATCAACTTGAAAAAGTGGCACCGAGTCGGTGCTTTTTTGTTTTAGAGCTAGAA
ATAGCAAGTTAAAATAAGGCTAGTCCGTTTTTAGCGCGTGCGCCAATTCTGCAGACA
AATGGCTCTAGAGGTACCCGTTACATAACTTACGGTAAATGGCCCGCCTGGCTGACC
GCCCAACGACCCCCGCCCATTGACGTCAATAGTAACGCCAATAGGGACTTTCCATTG
ACGTCAATGGGTGGAGTATTTACGGTAAACTGCCCACTTGGCAGTACATCAAGTGTA
TCATATGCCAAGTACGCCCCCTATTGACGTCAATGACGGTAAATGGCCCGCCTGGCA
TTGTGCCCAGTACATGACCTTATGGGACTTTCCTACTTGGCAGTACATCTACGTATTA
GTCATCGCTATTACCATGGTCGAGGTGAGCCCCACGTTCTGCTTCACTCTCCCCATCT
CCCCCCCCTCCCCACCCCCAATTTTGTATTTATTTATTTTTTAATTATTTTGTGCAGCG
ATGGGGGCGGGGGGGGGGGGGGGG-3'

5'-
ACMMMCGCMWWAMGATAMAAGGCTGTTAGAGAGATAATTGGAATTAATTTGACT
GTAAACACAAAGATATTAGTACAAAATACGTGACGTAGAAAGTAATAATTTCTTGG
GTAGTTTGCAGTTTTAAAATTATGTTTTAAAATGGACTATCATATGCTTACCGTAACT
TGAAAGTATTTCGATTTCTTGGCTTTATATATCTTGTGGAAAGGACGAAA**CACCGTG**
**ACTTACTCTCCAGAGTCA**GTTTTAGAGCTAGAAATAGCAAGTTAAAATAAGGCTAG
TCCGTTATCAACTTGAAAAAGTGGCACCGAGTCGGTGCTTTTTTGTTTTAGAGCTAG
AAATAGCAAGTTAAAATAAGGCTAGTCCGTTTTTAGCGCGTGCGCCAATTCTGCAGA
CAAATGGCTCTAGAGGTACCCGTTACATAACTTACGGTAAATGGCCCGCCTGGCTGA
CCGCCCAACGACCCCCGCCCATTGACGTCAATAGTAACGCCAATAGGGACTTTCCAT
TGACGTCAATGGGTGGAGTATTTACGGTAAACTGCCCACTTGGCAGTACATCAAGTG
TATCATATGCCAAGTACGCCCCCTATTGACGTCAATGACGGTAAATGGCCCGCCTGG
CATTGTGCCCAGTACATGACCTTATGGGACTTTCCTACTTGGCAGTACATCTACGTAT
TAGTCATCGCTATTACCATGGTCGAGGTGAGCCCCACGTTCTGCTTCACTCTCCCCAT
CTCCCCCCCCTCCCCACCCCCAATTTTGTATTTATTTATTTTTTAATTATTTTGTGCAG
CGATGGGGGCGGGGGGGGGGGGGGGGGSC-3'
